# Modeling CISH-KO TIL therapy: T-cell persistence and endogenous competition determine response in gastrointestinal cancer

**DOI:** 10.64898/2026.06.08.730755

**Authors:** Xuanming Zhang, Sarah Anderson, Kamran Kaveh, Matthew Johnson, David J. Odde, Beau Webber, Branden Moriarity, Emil Lou, Kevin Leder, Jasmine Foo

## Abstract

**Background:** Tumor-infiltrating lymphocyte (TIL) therapies have shown promise in murine models; however, early-phase clinical trials reveal substantial heterogeneity in treatment response across patients. Elucidating the biological drivers of this variability is critical for improving patient selection and therapeutic design.

**Methods:** We developed a mechanistic mathematical model of *Cish*-inactivated TIL therapy capturing interactions among tumor cells, endogenous TILs, and infused *CISH*-KO (*Cish* gene knockout) T-cells. The model was calibrated to longitudinal tumor size data from two murine studies and also fitted to clinical data from a first-in-human phase 1 trial enrolling 22 and treating 12 patients with metastatic gastrointestinal cancers (27 independently tracked tumor lesions). We used the calibrated model to identify patient-level determinants of response and to simulate combination and fractionation treatment strategies *in silico*.

**Findings:** Four mechanistically distinct parameters emerged as dominant determinants of therapeutic outcome: *CISH*-KO T-cell persistence (r = 0.827, p < 10^−4^) was the single strongest predictor of therapeutic failure; other major determinants included endogenous T-cell proliferation rate (r = 0.648, p < 10^−4^) , *CISH*-KO T-cell killing rate, and intrinsic tumor growth rate. In silico simulations predicted that the addition of anti–PD-1 checkpoint blockade to standard-dose *CISH*-KO TIL therapy achieved comparable or superior tumor control relative to dose doubling, while maintaining lower peak T-cell level. Additionally, fractionating the same total dose across multiple delayed infusions further improved predicted tumor control by prolonging CISH-KO T-cell persistence, again without raising peak effector burden.

**Interpretation:** The efficacy of current *CISH*-KO TIL protocols is influenced not only by the cytotoxic potency of infused T-cells, but also by their persistence within the tumor microenvironment and competitive pressure from the reconstituting host immune compartment. These findings support strategies that enhance the functional efficiency and prolong the availability of infused T-cells—such as concurrent PD-1 blockade or fractionated dosing of the same total cell product—rather than dose escalation alone and identify *CISH*-KO T-cell persistence and endogenous immune competition as actionable targets for improving TIL-based immunotherapies.

## Introduction

T-cells play a critical role in cancer progression and prevention, as cytotoxic T-cells are able to infiltrate tumors and kill cancer cells (1,2). Conversely, this activity is counterbalanced through immunosuppressive regulatory T-cells and macrophages (3,4). In a variety of cancers, the density, distribution, and functional activity of tumor-infiltrating T-cells correlates with patient prognosis, with higher levels of T-cell infiltration and activity associated with better survival (5–8). These observations have driven the development of therapies aimed at mobilizing the immune system to target cancer cells (9,10). One such approach is autologous tumor-infiltrating lymphocyte (TIL) therapy, in which lymphocytes isolated from patient tumor samples are expanded *ex vivo* and reinfused. Although objective responses have been reported in select patients and cancer types, the efficacy of unedited TIL therapies in solid tumors remains limited. Key barriers to efficacy may include T-cell exhaustion, immunosuppressive signaling within the tumor microenvironment, and on-target/off-tumor toxicity (11–14) . Overcoming these challenges is therefore essential for improving the durability and breadth of TIL-based immunotherapies.

Immune activation, differentiation, and cytotoxic function are regulated by a complex network of signals within the tumor microenvironment and systemic immune milieu. Among these regulators, the suppressors of the cytokine signaling (SOCS) family—comprising Socs1–7 and *Cish*—play a critical role in modulating immune responses. SOCS proteins negatively regulate cytokine and receptor signaling, and their manipulation has been shown to enhance antitumor immunity (15,16). *Cish* functions as an intracellular immune checkpoint in T-cells and natural killer cells, with expression induced following T-cell receptor stimulation or cytokine exposure, including IL-2 (17,18). Elevated *Cish* expression has been observed in TILs, where it suppresses antitumor activity (19). Genetic deletion of *Cish* in CD8^+^ T-cells enhances expansion, functional avidity, and polycytokine secretion, and induces a metabolic and activation program characterized by upregulation of multiple activation markers, including PD-1 (19). These findings suggest that *Cish* disruption may improve the efficacy of TIL immunotherapy, a notion that has been supported by preclinical studies demonstrating the therapeutic potential of *Cish*-deficient TILs. In murine B16 melanoma models, adoptive transfer of *Cish−*KO CD8^+^ T-cells resulted in complete tumor eradication, in contrast to wild-type T-cell transfer or no treatment (16). Subsequent studies showed that, in the absence of exogenous IL-2, *Cish*-KO T-cells slowed but did not fully arrest tumor growth; however, combination with anti–PD-1 therapy restored complete tumor clearance (19). These promising results motivated a first-in-human clinical trial in which primary human T-cells were engineered via CRISPR/Cas9 to inactivate *Cish* and administered alongside IL-2 to patients with gastrointestinal cancer (20). The trial demonstrated safety and produced stable disease in 50% of patients, as well as a durable complete response in 1 of the 12 end stage patients treated – demonstrating the therapeutic potential of gene-edit TIL therapy. However, clinical responses were heterogeneous and substantially more variable than those observed in murine models (16,19), suggesting that therapeutic efficacy is governed by complex biological determinants that are not fully captured by preclinical systems.

The complexity of the tumor–immune interactions underlying this variability spans tumor-intrinsic growth dynamics, immune competence, T-cell exhaustion, and treatment protocol. Descriptive analyses of trial data can identify factors correlating with response, but are often limited in providing mechanistic insights into the underlying sources of variability. Elucidating these mechanistic contributions motivates the use of mathematical modeling to disentangle and quantify the relative contributions of these factors (21–23).

Specifically, this requires a mathematical modeling framework that is capable of representing complex tumor and immune interactions within the tumor microenvironment, fitting to clinical longitudinal data, and exploring in silico perturbations to investigate alternative treatment and parameter scenarios. In this study, we develop and calibrate a mechanistic ordinary differential equation (ODE) model to investigate the determinants of response to *CISH-*KO TIL therapy. Using tumor response data from murine studies, we inform and validate the model structure, and subsequently apply it to interpret heterogeneous outcomes observed in a human clinical trial (20). We leverage this framework to identify key factors differentiating responders from non-responders, to explore mechanistic explanations for discrepancies between murine and human efficacy, and to perform simulated clinical trials evaluating alternative therapeutic strategies.

Our analysis identifies four mechanistically distinct determinants of therapeutic outcome: the intrinsic tumor growth rate, the *CISH*-KO T-cell death rate, the endogenous T-cell proliferation rate, and the *CISH*-KO T-cell killing rate. While the roles of tumor growth and cytotoxic potency are intuitive, our results reveal that limited T-cell persistence was the strongest determinant of response, and that competitive displacement by functionally inferior endogenous T-cells is also an important factor. These findings indicate that the efficacy of current *CISH*-KO TILs is constrained not only by the potency of the infused cells, but also by their survival within the tumor microenvironment and competition from the reconstituting host immune compartment. Guided by preclinical evidence of synergy between *Cish* deletion and PD-1 blockade, we further evaluate combination therapy and fractionated dosing strategies in silico. Our simulations predict that combination therapy achieves superior tumor control relative to monotherapy or dose escalation, and that fractionating the same total cell product across delayed infusions further improves outcomes by prolonging CISH-KO T-cell persistence, both without increasing peak T-cell exposure, suggesting a pathway to improved efficacy while maintaining a favorable safety profile.

## Methods

### Data Sources

*Murine data.* Data were digitized from *in vivo* experiments reported in (16,19). In both studies, melanoma/melanocyte-specific pmel-1 T-cells, with or without *Cish*, were adoptively transferred into established B16 melanoma–bearing C57BL/6 mice. Longitudinal tumor size measurements under the various treatment conditions were extracted from the published figures using WebPlotDigitizer.

In the 2015 study, mice bearing established B16 melanoma tumors (tumor area approximately 40 mm^2^ at day 0) were assigned to three groups: untreated controls, mice receiving adoptively transferred wild-type T-cells, and mice receiving *Cish*-KO T-cells. Tumor area was measured longitudinally over a 32-day observation window.

The 2022 study extended this design to six groups by further stratifying mice based on the presence or absence of anti–PD-1 treatment. Mice bearing established tumors (tumor area approximately 25 mm^2^ at day 0) were randomized into: (1) no T-cells, isotype control; (2) no T-cells + anti–PD-1; (3) wild-type T-cells, isotype control; (4) wild-type T-cells + anti–PD-1; (5) *Cish*-KO T-cells, isotype control; (6) *Cish*-KO T-cells + anti–PD-1. Tumor area was measured longitudinally over a 42-day observation window. The inclusion of anti–PD-1 arms in the 2022 study enabled characterization of the interaction between *Cish*-KO and immune checkpoint blockade.

*Clinical trial data.* Patient data were obtained from a first-in-human, single-centre, phase I clinical trial (24) enrolling patients aged 18–70 years with metastatic gastrointestinal epithelial cancers. Tumor-infiltrating lymphocytes (TILs) were isolated from tumor biopsies, expanded based on neoantigen reactivity, subjected to CRISPR–Cas9–mediated *CISH*-KO, and intravenously infused into 12 patients following non-myeloablative lymphodepleting chemotherapy. Dose assignment followed a phase I dose-escalation design, with escalation proceeding until dose-limiting toxicity or the highest dose level (dose level 5) was reached without dose-limiting toxicity.

Three types of longitudinal data were available for model calibration:

**Tumor size measurements.** Tumor burden was assessed approximately monthly by imaging. Longitudinal measurements were available for a total of 27 tumor lesions across primary and metastatic sites, with individual patients harboring between 1 and 5 independently tracked lesions. With the exception of one patient (UMN022), tumors were either stable or increasing in size over the observation period; patient UMN022 exhibited a clear trend of tumor regression.
**Circulating *CISH*-KO T-cell frequency.** The ratio of *CISH-*KO to wild-type T-cells in peripheral blood was quantified by flow cytometry from peripheral blood mononuclear cell (PBMC) samples collected at multiple time points post-infusion. Because direct measurement of this ratio among tumor-infiltrating lymphocytes is not feasible *in vivo*, we used the circulating ratio as a proxy for the corresponding ratio within the tumor microenvironment. This assumption is motivated by studies demonstrating that the persistence and expansion of adoptively transferred T-cell clonotypes in peripheral blood correlate with clinical tumor regression (25,26), suggesting that circulating T-cell dynamics are informative about therapeutic activity at the tumor site. However, differential trafficking and local expansion may cause the intratumoral ratio to diverge from the peripheral measurement, particularly at later time points; we discuss this limitation further in the Discussion.
**Infused T-cell dose.** Each patient received a single intravenous infusion of *CISH*-KO T-cells at day 0. The total number of engineered cells infused per patient was recorded, with doses typically on the order of 10^10^ cells.

### Mathematical Model

To interpret results from mouse and human trials of *CISH*-KO TIL therapy, we developed a mechanistic ODE model capturing the key interactions governing tumor response to adoptive T-cell therapy (Figure 1 - model schematic, Supplementary Material - equations and rationale). We began with a biologically comprehensive model incorporating all plausible interactions among tumor cells and the three T-cell populations, and subsequently applied systematic identifiability analysis to determine which parameters can be resolved from the available data. Non-identifiable parameters were fixed at literature-informed values. This approach ensures that model simplification is driven by the information content of the data rather than by *a priori* assumptions about which biological processes are negligible, and avoids the risk of prematurely excluding mechanisms that may become identifiable with richer datasets.

**Figure 1.**
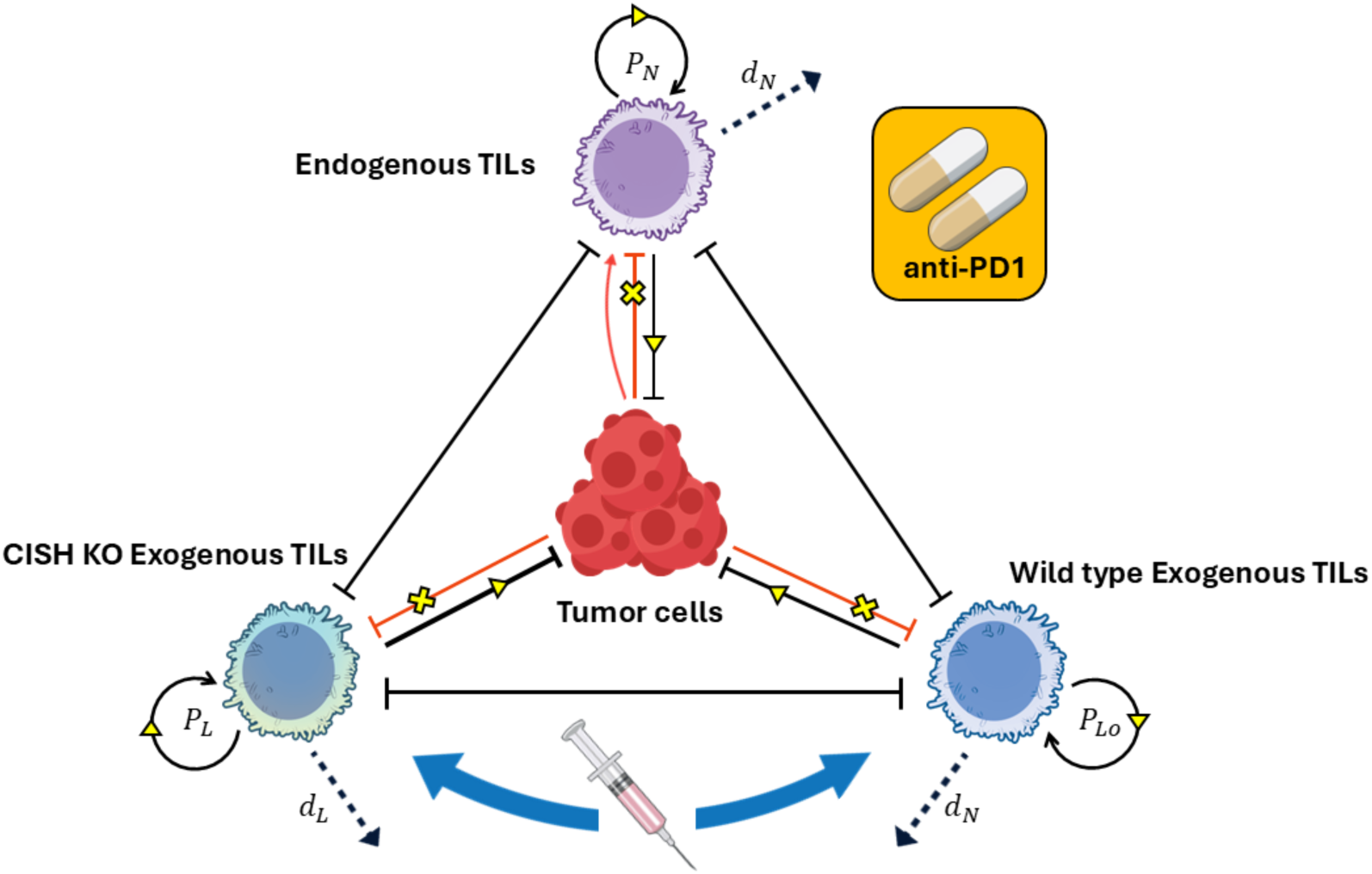
: Schematic of tumor–immune interactions under *CISH-KO* TIL infusion and anti–PD-1 therapy. The diagram illustrates the interactions between tumor cells (center) and three T-cell populations: endogenous TILs (top), exogenous *CISH-KO* TILs (left), and exogenous wild-type TILs (right). Tumor growth is suppressed through cytotoxic tumor–T-cells interactions, which also promote the recruitment and expansion of endogenous TILs. The relative tumor-killing efficacy satisfies *CISH-KO* TILs > wild-type TILs > endogenous TILs. Competition among T-cell populations arises from a shared carrying capacity of the immune compartment, leading to proliferation suppression. Yellow markers denote the therapeutic effects of anti–PD-1 treatment, including enhanced proliferation, T-cell-mediated tumor killing and inhibition of tumor-induced T-cell exhaustion.

The model represents the tumor microenvironment through four interacting components: tumor cells (T) and three T-cell populations—endogenous tumor-infiltrating lymphocytes (TILs; N), exogenously infused *CISH-*inactivated TILs (L), and exogenous wild-type TILs (L_o_). Endogenous TILs represent the native immune response with limited tumor-specific recognition, whereas the two exogenous populations correspond to adoptively transferred T-cells. Among these, *CISH*-KO TILs exhibit enhanced cytotoxicity due to loss of *CISH*-KO, while wild-type TILs represent baseline adoptive cell therapy without genetic modification.

The core model dynamics are as follows. Tumor cells proliferate exponentially and are eliminated through cytotoxic interactions with all three T-cell populations. Each T-cell population undergoes natural death and tumor-induced exhaustion, and proliferates subject to logistic competition for a shared immune carrying capacity. This logistic formulation is a standard framework for modeling resource-limited competition among T-cell populations, originally developed in the context of T-cell repertoire dynamics[19], and reflects competition for common limiting resources including cytokines, antigen-presenting cell contacts, and spatial niches within the tumor microenvironment. Endogenous TILs are additionally recruited in response to tumor burden via a saturation-dependent accumulation term. After intravenous infusion, transferred T cells do not immediately accumulate within the tumor, as some cells may be cleared, remain in circulation, or distribute to non-tumor tissues. We therefore represent the infused cell dose as a transient in vivo reservoir, from which cells gradually infiltrate the tumor while some are lost before engraftment. Anti–PD-1 therapy, when present, enhances T-cell proliferation and cytotoxicity while attenuating tumor-induced exhaustion, with separate response functions for wild-type and *CISH*-KO T-cells to reflect their distinct susceptibilities to PD-1 blockade(19).

A schematic overview is shown in Figure 1 and the full system of equations, parameter definitions, and model selection rationale are provided in the Supplementary Material.

### Model Calibration and Goodness of Fit

Our modeling calibration strategy follows a deliberate parameter elimination approach to guarantee the identifiability of chosen parameters. The model was calibrated separately for both murine and clinical datasets. In both cases, model parameters were estimated by minimizing the discrepancy between simulated and observed tumor trajectories. For the murine data, the objective function employed scale-invariant relative-error weighting across treatment arms to normalize residuals relative to observed maximum tumor size, ensuring that arms with effective therapy (and consequently lower tumor volumes) contributed fairly to the fit. For the clinical data, a scale-invariant relative-error formulation was used, eliminating the need for manual weighting across lesions of different sizes. The detailed fitting procedures, objective functions, optimization algorithms, parameter bounds, and identifiability analyses are described in the Supplementary Material.

### Mouse model calibration

The model was simultaneously fitted to longitudinal tumor size measurements from all treatment arms within each mouse study. The 2015 study(16) contributed three treatment conditions (no treatment, wild-type T-cells, and engineered *Cish*-KO T-cells), while the 2022 study(19) contributed six conditions including anti–PD-1 combination arms. As shown in Figure 2, the calibrated model accurately reproduces the differential tumor growth trajectories across all treatment arms, capturing both the transient tumor expansion prior to immune-mediated suppression and the sustained regression under combination therapy. Observed versus predicted tumor sizes yielded R^2^ = 0.928 for the 2015 study and R^2^ = 0.935 for the 2022 study (Figure 2c,d).

**Figure 2.**
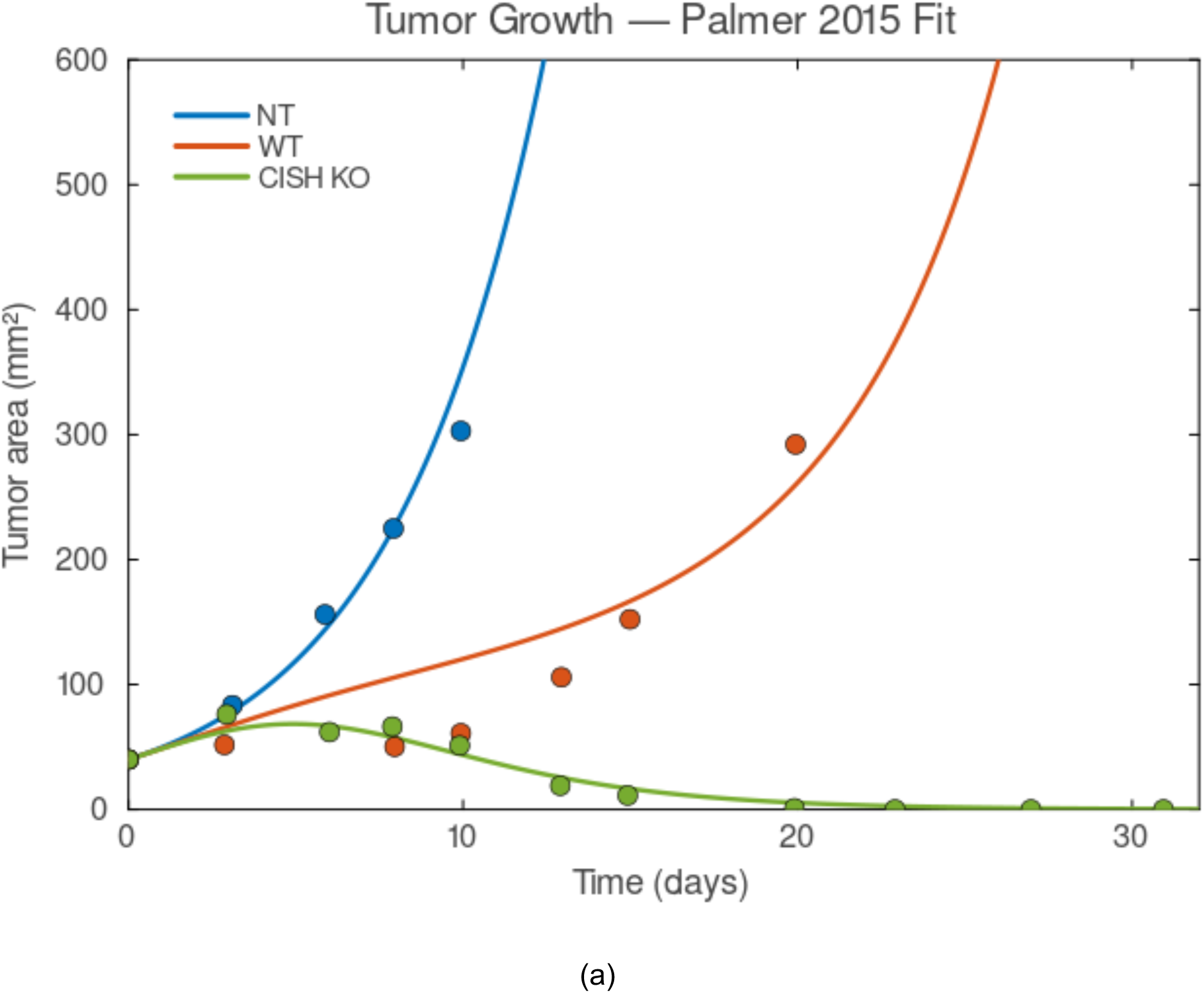
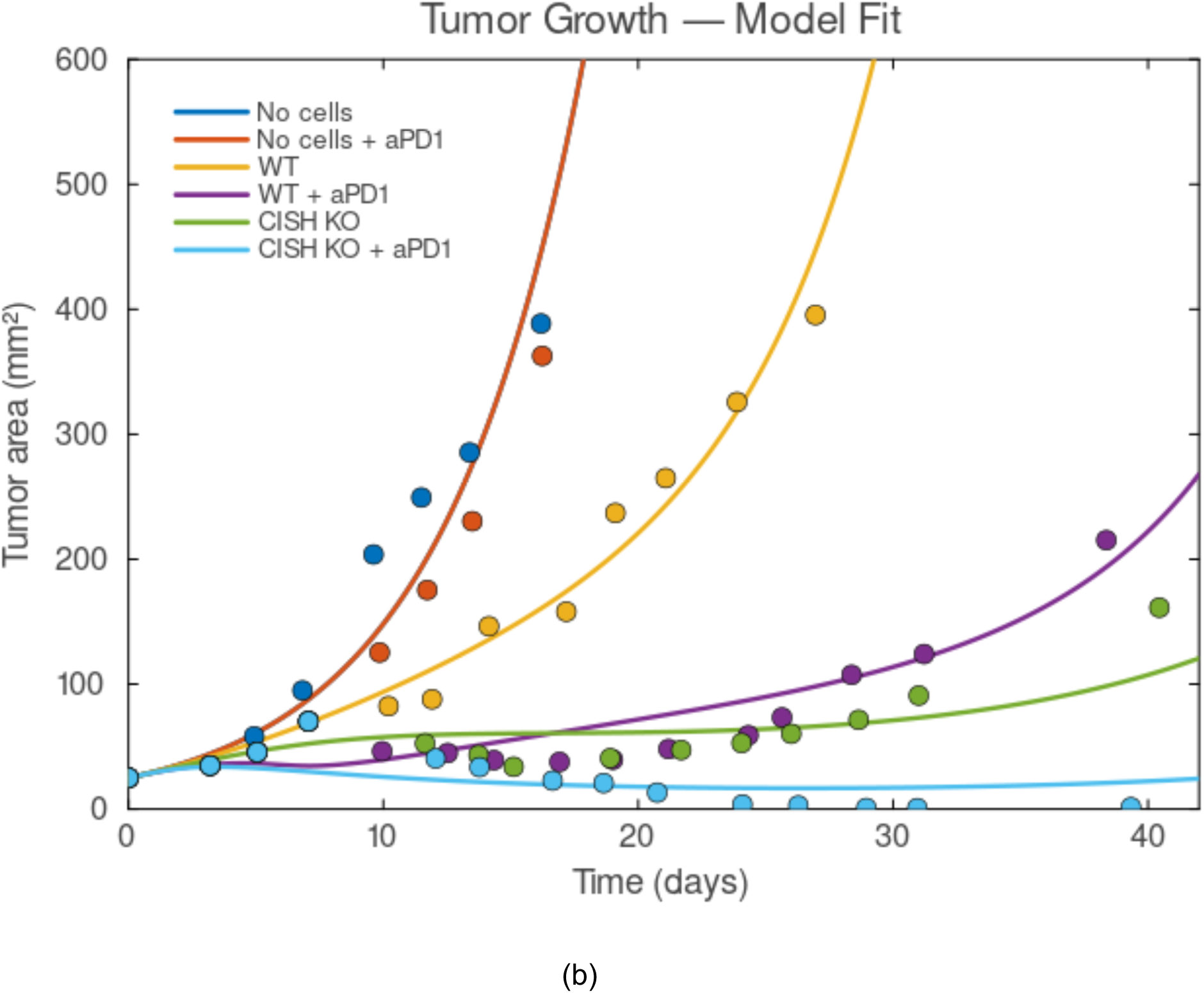
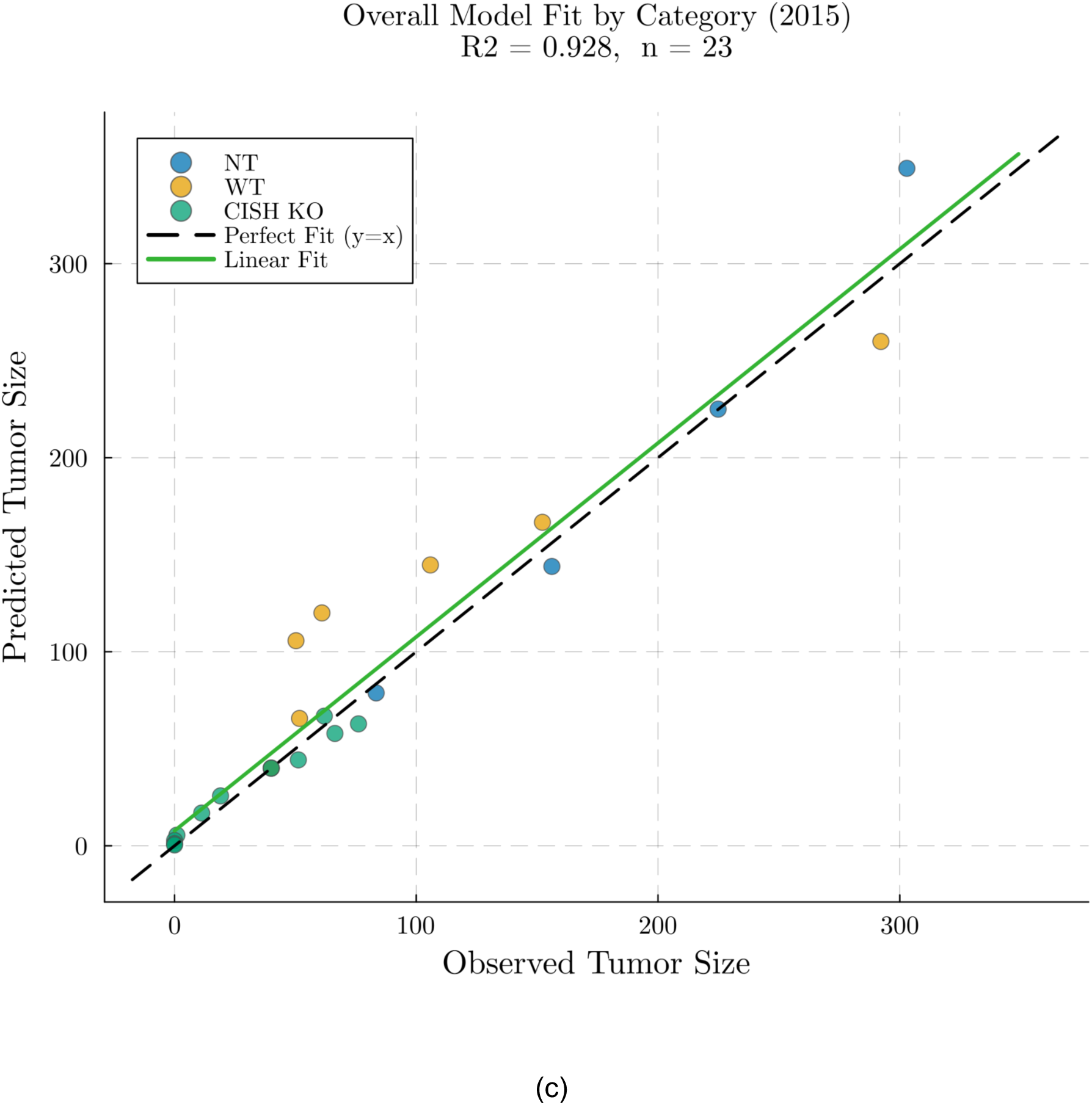
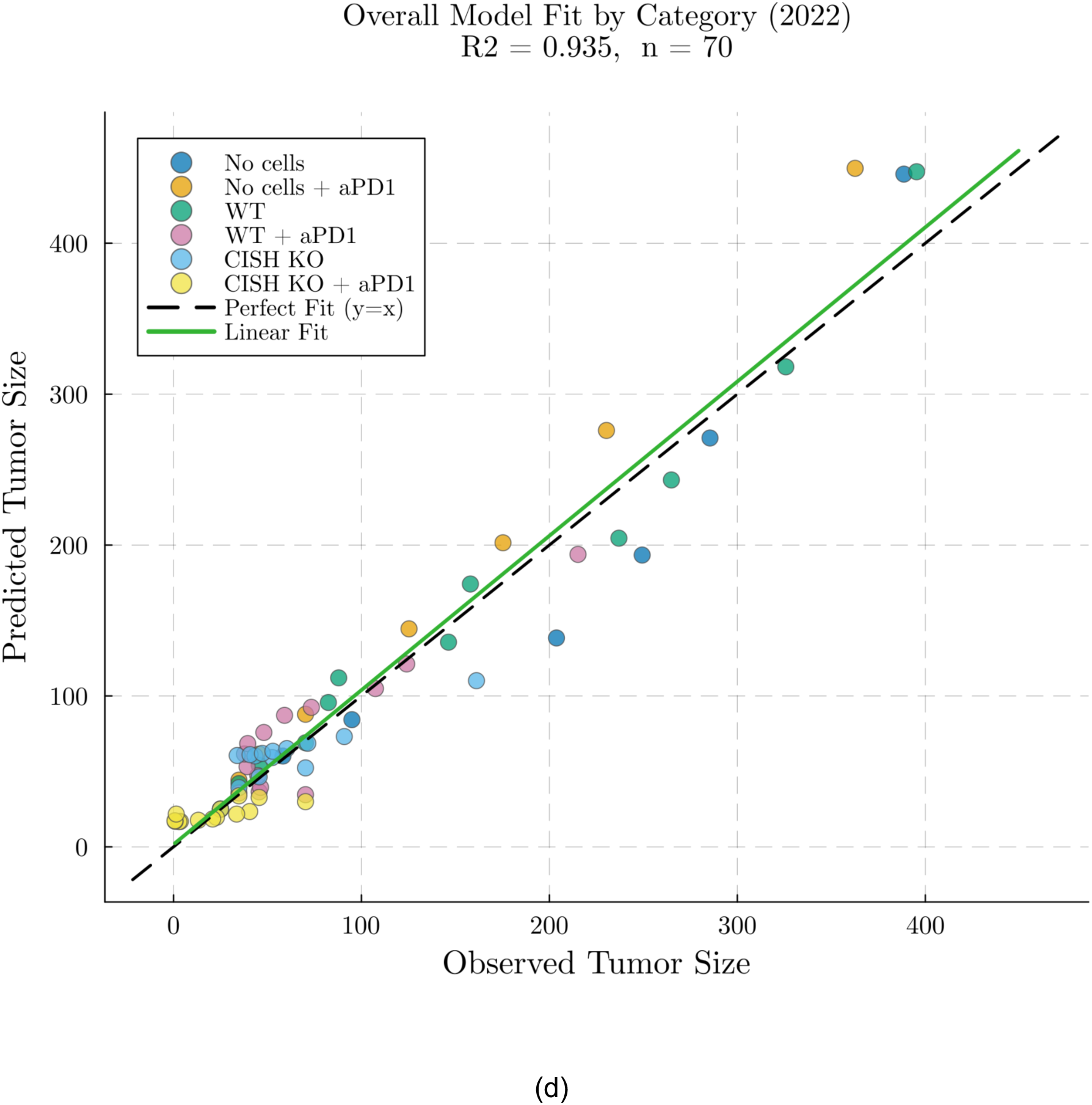
: **Model calibration against *in vivo* murine tumor growth data. (a, b)** Solid lines denote model simulations obtained by fitting the ODE system to longitudinal tumor size measurements (dots). (a) Fits to data from Palmer et al. [13] demonstrate the model’s ability to capture differential therapeutic efficacy, with *Cish*-KO T-cells (yellow) producing stronger tumor suppression than wild-type (WT) T-cells (red) and no-treatment controls (NT, blue). (b) Fits to data from (19) incorporating anti–PD-1 checkpoint blockade. The model reproduces the synergistic inhibition of tumor growth, realizing sustained tumor regression under combined *Cish*-KO and anti–PD-1 therapy (light blue), outperforming WT + anti–PD-1 (purple) and single-agent treatments. **(c, d)** Observed vs. predicted tumor size for the 2015 (R^2^ = 0.928, n = 23) and 2022 (R^2^ = 0.935, n = 70) experiments. Each point represents a single time-point observation color-coded by treatment arm. The dashed black line is the identity (y = x); the solid green line is the ordinary least-squares regression.

### Clinical model calibration

For the clinical dataset, we employed a hierarchical optimization framework to account for both inter-patient and intra-patient variability. A subset of model parameters was allowed to vary at the patient level to capture differences in T-cell survival, cytotoxic efficacy, proliferation, and infiltration efficiency, while the tumor intrinsic growth rate was fitted independently for each lesion. The selection of patient-level parameters was guided by an iterative procedure combining pairwise correlation analysis and profile likelihood assessment to ensure identifiability. As a result, parameters governing individual endogenous T-cell kinetics were non-identifiable from the clinical data and fixed at nominal values from mouse model fitting, whereas parameters controlling *CISH*-KO T-cell dynamics and the net endogenous reconstitution rate were retained as they govern mechanistically distinct processes with distinguishable signatures in the observed data (see Supplementary Material, Section S2.4 for details).

The fitted model showed strong agreement with clinical observations. As shown in Figure 3a, model-predicted tumor volumes were highly correlated with observed values across all 27 lesions (R^2^ = 0.996, RMSE = 1.62). The model also captured the longitudinal pattern of circulating *CISH*-KO T-cell ratios (Figure 3b), which were used as a proxy for intratumoral composition. Deviations were most apparent at very low ratio values, where the continuous ODE framework predicts asymptotically small populations while experimental measurements are bounded by assay detection limits. Individual patient fits are shown in Supplementary Figures S7–S9.

**Figure 3.**
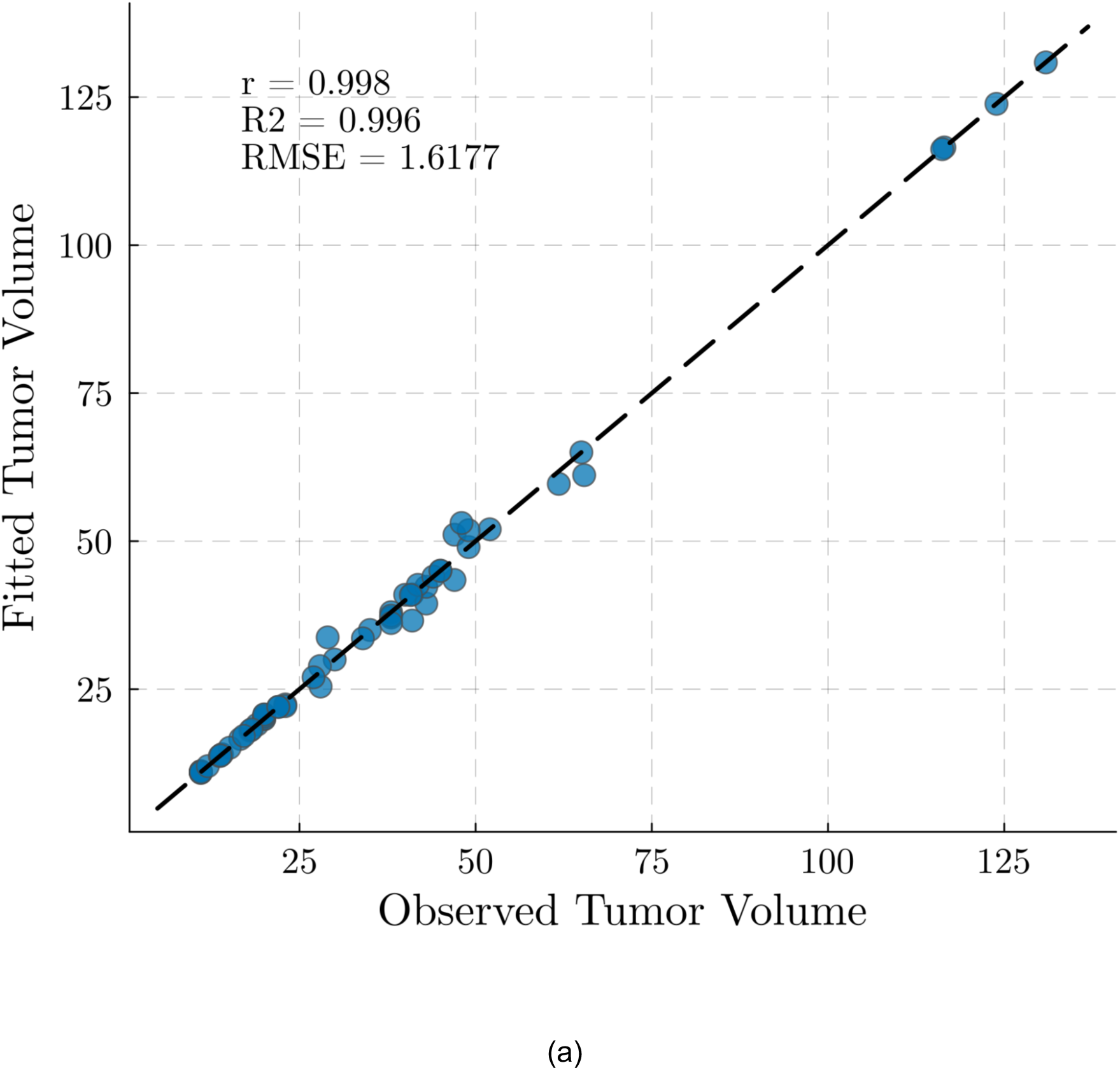
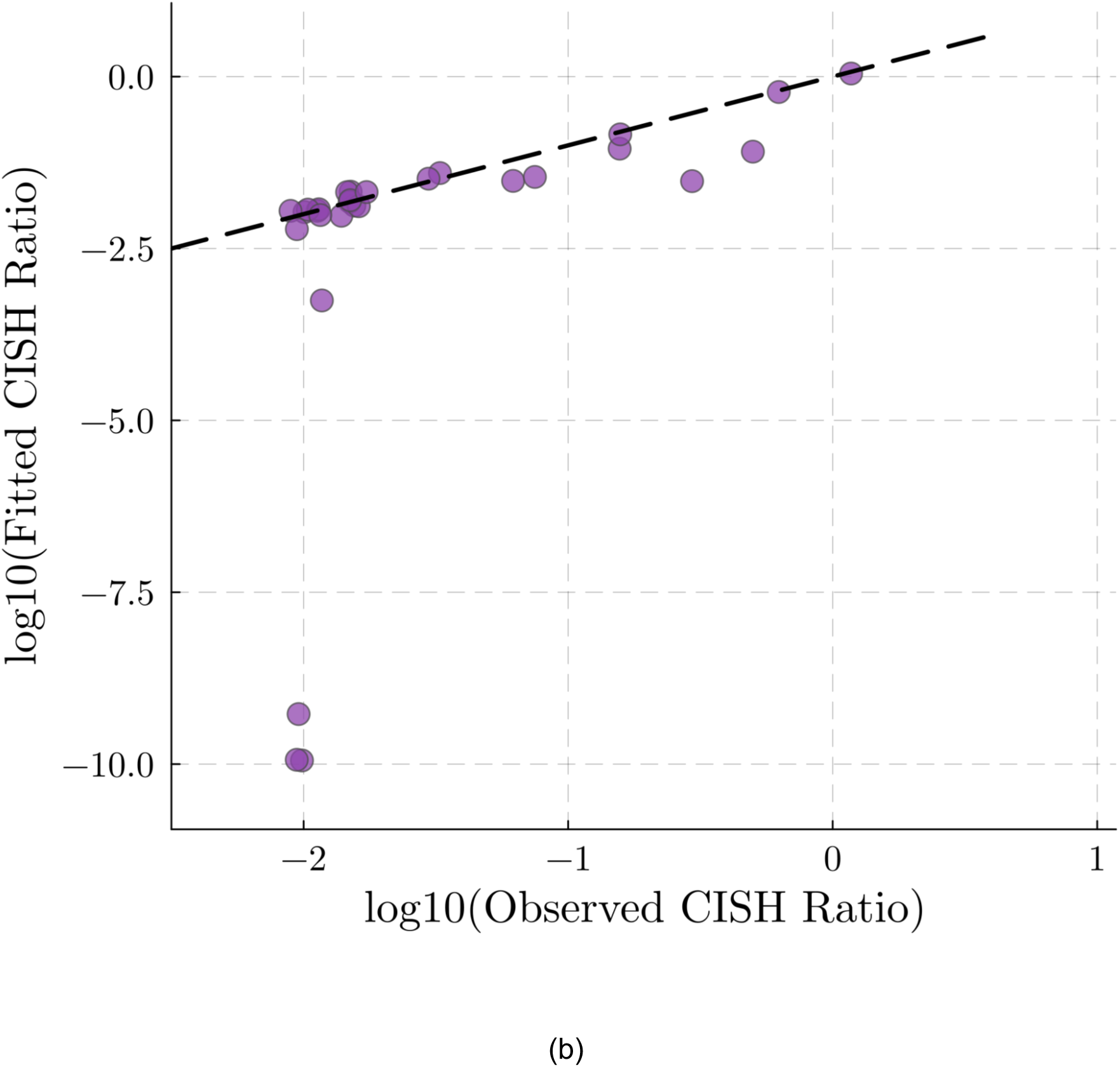
: **Model goodness-of-fit diagnostics for clinical data. (a)** Comparison of fitted versus observed tumor volumes across all lesions demonstrates high predictive accuracy (R^2^ = 0.996, RMSE = 1.62, r = 0.998 representing Pearson’s correlation). **(b)** Log-transformed comparison of fitted versus observed circulating *CISH-KO* T-cell ratios. The model aligns well with observed ratios in active disease states, while deviations at very low values reflect differences between the continuous ODE framework and experimental measurement limits. Specifically, the model drives strongly suppressed populations toward asymptotically small values (fitted ratios < 10^−6^), whereas experimental measurements are constrained by a noise floor (approximately 10^−2^), leading to apparent divergence in this regime.

## Results

### Parameter fitting reveals conserved T-cell cytotoxicity hierarchy in mice

Calibration of the model with murine data (16,19) yielded fitted parameter values reported in Supplementary Table 2. Because both experiments employed the B16 melanoma model in C57BL/6 mice, most physiological parameters were consistent across fits. In particular, the tumor killing rates k_L_ and k_Lo_ are comparable between the two studies, as expected given that both employed the same TCR-engineered T-cell product in the same tumor model. The primary differences between fits were in the death rate d_L_ of *CISH*-KO T-cells, which can be attributed to the use of IL-2 in the 2015 study (which is not explicitly modeled) as well as experiment-to-experiment variability.

Across both datasets, the fitted cytotoxicity parameters reveal a clear hierarchy of killing efficacy, with k_L_ > k_Lo_ > k_N_. The markedly lower killing rate of endogenous T-cells (k_N_) relative to infused wild-type and *CISH*-KO T-cells suggests limited functional avidity of native TCR repertoires compared to *ex vivo* selected and expanded TIL products. This finding provides a mechanistic explanation for the limited efficacy of anti–PD-1 monotherapy alone: while checkpoint blockade can enhance T-cells cytotoxicity, it cannot compensate for insufficient baseline tumor recognition (Figure 4a).

**Figure 4.**
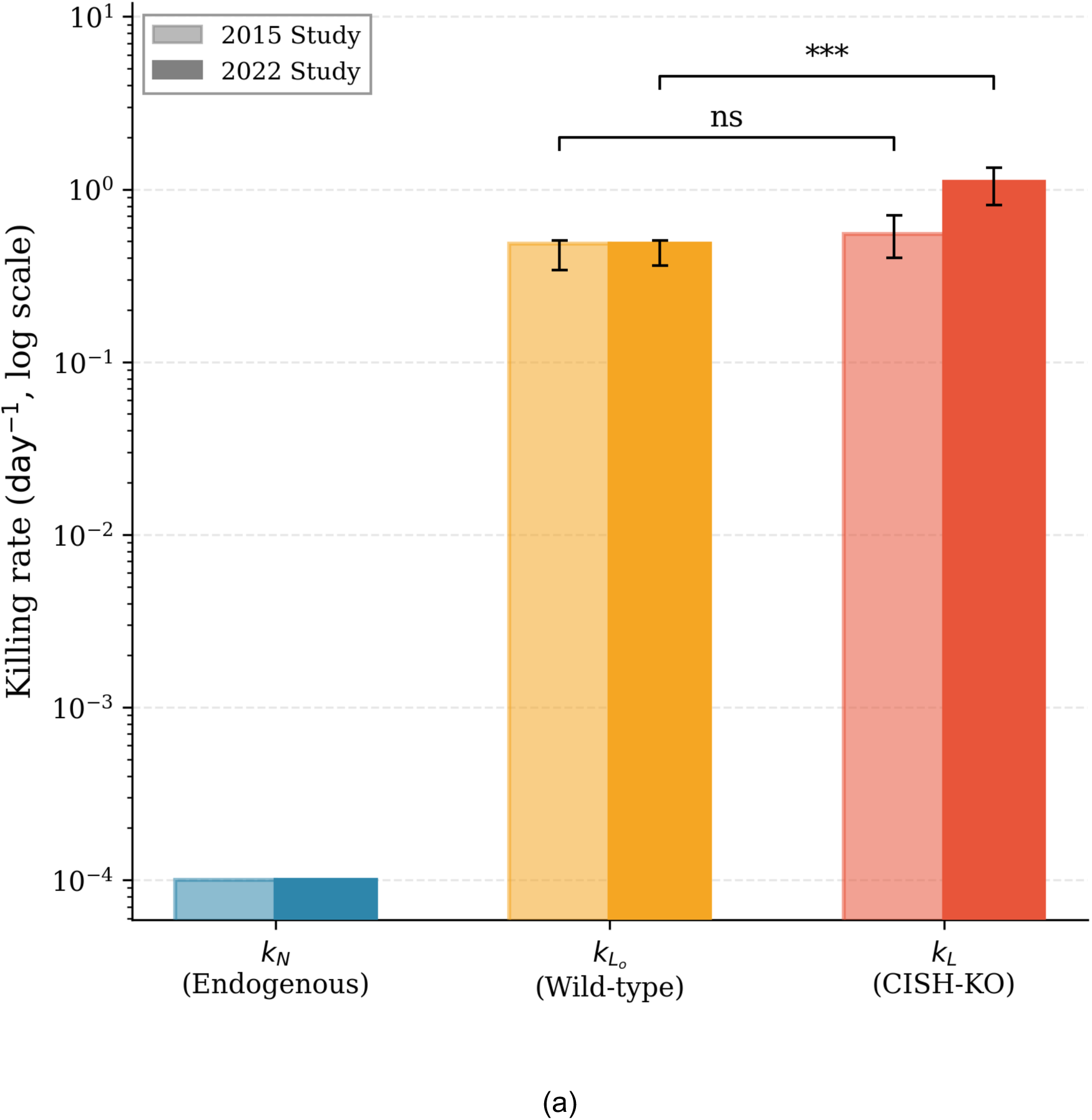
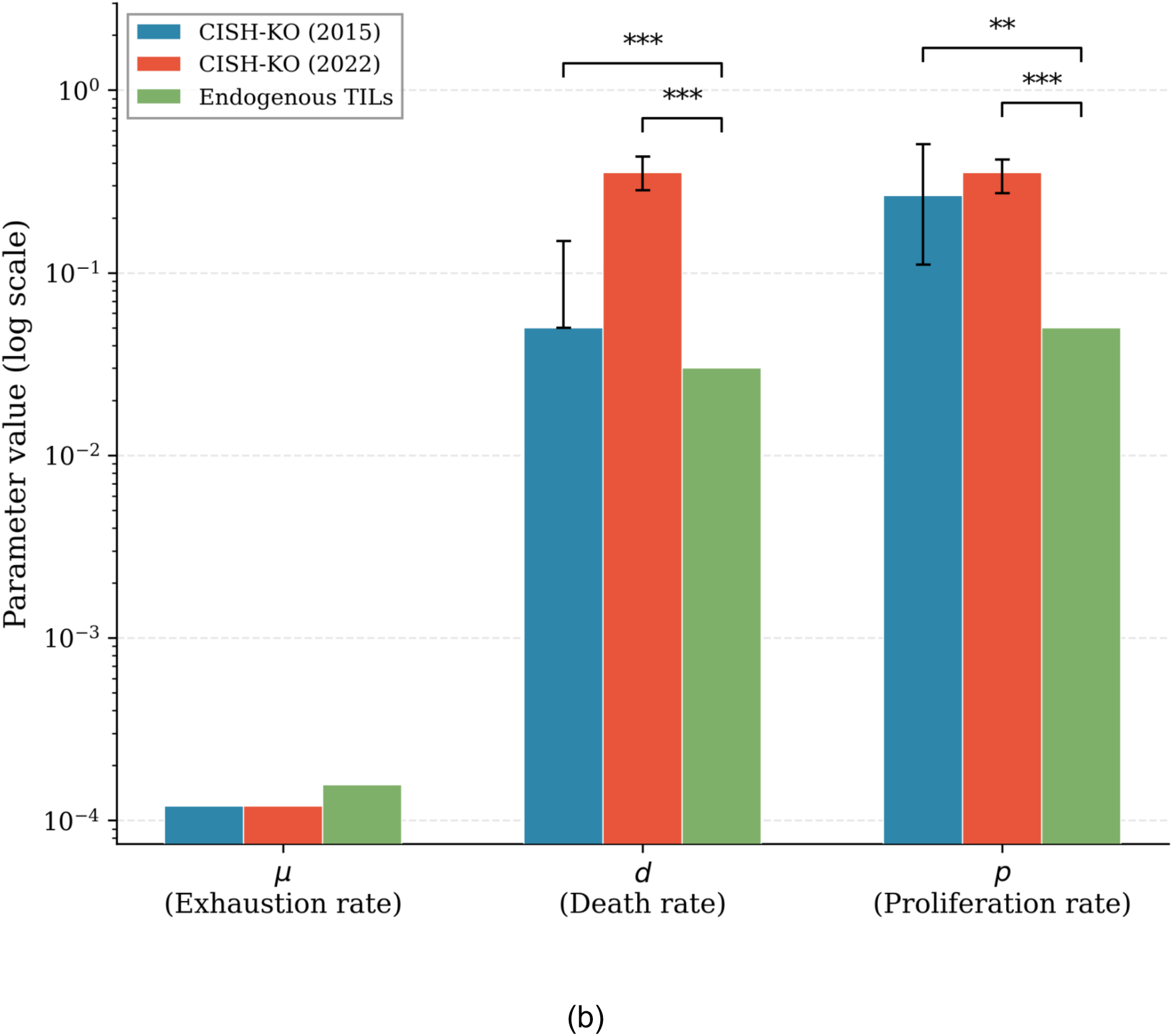
: **Fitted parameters reveal conserved cytotoxicity hierarchy across murine studies (a)** Tumor killing rates across the three T-cell populations for each study. The ordering k_L_ > k_Lo_ > k_N_ is preserved in both experiments, confirming the enhanced per-cell cytotoxicity of *CISH-KO* T-cells relative to wild-type and endogenous TILs. Lighter and darker shading denote the 2015 and 2022 studies, respectively. **(b)** Comparison of exhaustion rate (μ), natural death rate (d), and proliferation rate (p) for *CISH-KO* T-cells (2015 and 2022 fits) versus endogenous TILs (log scale). The key comparison is between *CISH-KO* and endogenous populations: endogenous TILs have slightly lower proliferation (p_N_), while *CISH-KO* T-cells exhibit higher death rates than the endogenous compartment.

The fitted proliferation and death rates further reflect known biological trade-offs. *CISH*-KO T-cells exhibit elevated proliferation rates p_L_ and death rates d_L_, aligning with observations that enhanced activation can compromise long-term persistence. Collectively, these findings indicate that *CISH* disruption primarily enhances per-cell cytotoxic potency rather than durability, a distinction that is critical for interpreting downstream therapeutic limitations (Figure 4b).

### Clinical parameter fitting reveals reduced *CISH*-KO T-cell cytotoxicity and competitive capacity in human patients

Calibrating the model with *CISH*-KO TIL phase I trial data (24) resulted in fitted parameter values for each patient, which are reported in Supplementary Table 4. Comparison of mouse and clinical fitting parameters reveals key species-specific differences. The *CISH*-KO T-cell killing rate (k_L_) was substantially lower in clinical patients compared to mouse models, reflecting reduced per-cell cytotoxic potency in the human setting.

Additionally, the ratio of *CISH*-KO to endogenous T-cell proliferation rates (p_L_/p_N_) was much higher in mouse experiments, indicating that infused *CISH*-KO T-cells outcompete endogenous populations more effectively in mouse models. These differences are illustrated in Figure 5a. Other endogenous T-cell kinetic parameters were fixed at nominal mouse-derived values because they could not be resolved from the clinical data, highlighting the limited information content of human observations regarding native immune reconstitution.

**Figure 5.**
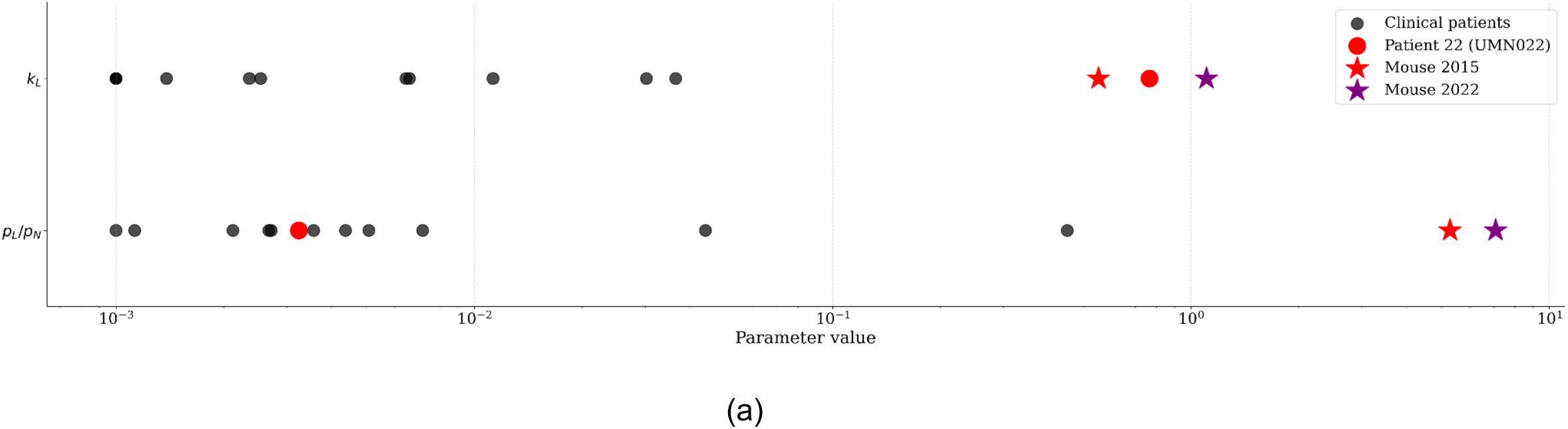
:Fitted parameters identify reduced killing and proliferative capability of CISH-KO T-cells in clinical translation relative to mouse models. (a) Comparison of key model parameters fitted to murine versus clinical data, highlighting the reduced killing rate (k_L_), proliferation advantage (p_L_/p_N_), and persistence of *CISH-KO* T-cells in human patients relative to IL-2-supported mouse models.

These differences have important biological implications. The reduced p_L_/p_N_ ratio in clinical patients indicates that infused CISH-KO T-cells do not outcompete endogenous populations as effectively as in mouse models, suggesting diminished competitive expansion in the human setting. The lower k_L_ in clinical patients corresponds to the poorer therapeutic performance observed in most cases, with the notable exception of patient UMN022, whose k_L_ values were comparable to those in mouse experiments and associated with durable tumor regression. Potential mechanistic explanations for these species-level differences, including the role of the Rapid Expansion Protocol in clinical manufacturing, are considered in the Discussion. In the next section, we quantify how variability in these identifiable parameters shapes patient-specific therapeutic outcomes.

### *CISH*-KO T-cell persistence and endogenous competition emerge as dominant determinants of response

Clinical responses to *CISH*-KO TIL therapy exhibited substantial heterogeneity across patients. While most tumor lesions remained stable or increased in size over the treatment period, one patient (UMN022) displayed pronounced tumor suppression followed by elimination of previously visualized metastatic disease. Elucidating the biological mechanisms underlying this variability is essential for understanding the limitations of current *CISH*-KO TIL therapy and for identifying strategies to improve its efficacy through further bioengineering.

To quantify therapeutic outcomes, we focused on four summary indices derived from the fitted model: (i) the tumor ratio, defined as the final tumor size normalized by the initial size; (ii) the normalized tumor exposure, measured as the tumor area under the curve (AUC) divided by the initial tumor size; (iii) the maximum *CISH*- KO T-cell abundance; and (iv) the *CISH*-KO T-cell AUC, representing cumulative T-cell exposure over the treatment period.

To identify key drivers of inter-patient variability within the trial responses, we examined associations between therapeutic outcome metrics and fitted model parameters across all 27 tumor lesions. Scatter plots in Figure 6a–d reveal that four parameters are strongly associated with treatment response: the intrinsic tumor growth rate γ, the *CISH*-KO T-cell death rate d_L_, the endogenous T-cell proliferation rate p_N_, and the *CISH*-KO T-cell killing rate k_L_. Specifically, higher values of γ (r = 0.653, p < 10^−4^) and d_L_ (r = 0.827, p < 10^−4^) are associated with elevated tumor ratios, while higher p_N_ (r = 0.648, p < 10^−4^) is associated with increased normalized tumor exposure. Conversely, higher k_L_ (r = −0.426, p = 0.0096) is associated with reduced tumor exposure.

**Figure 6.**
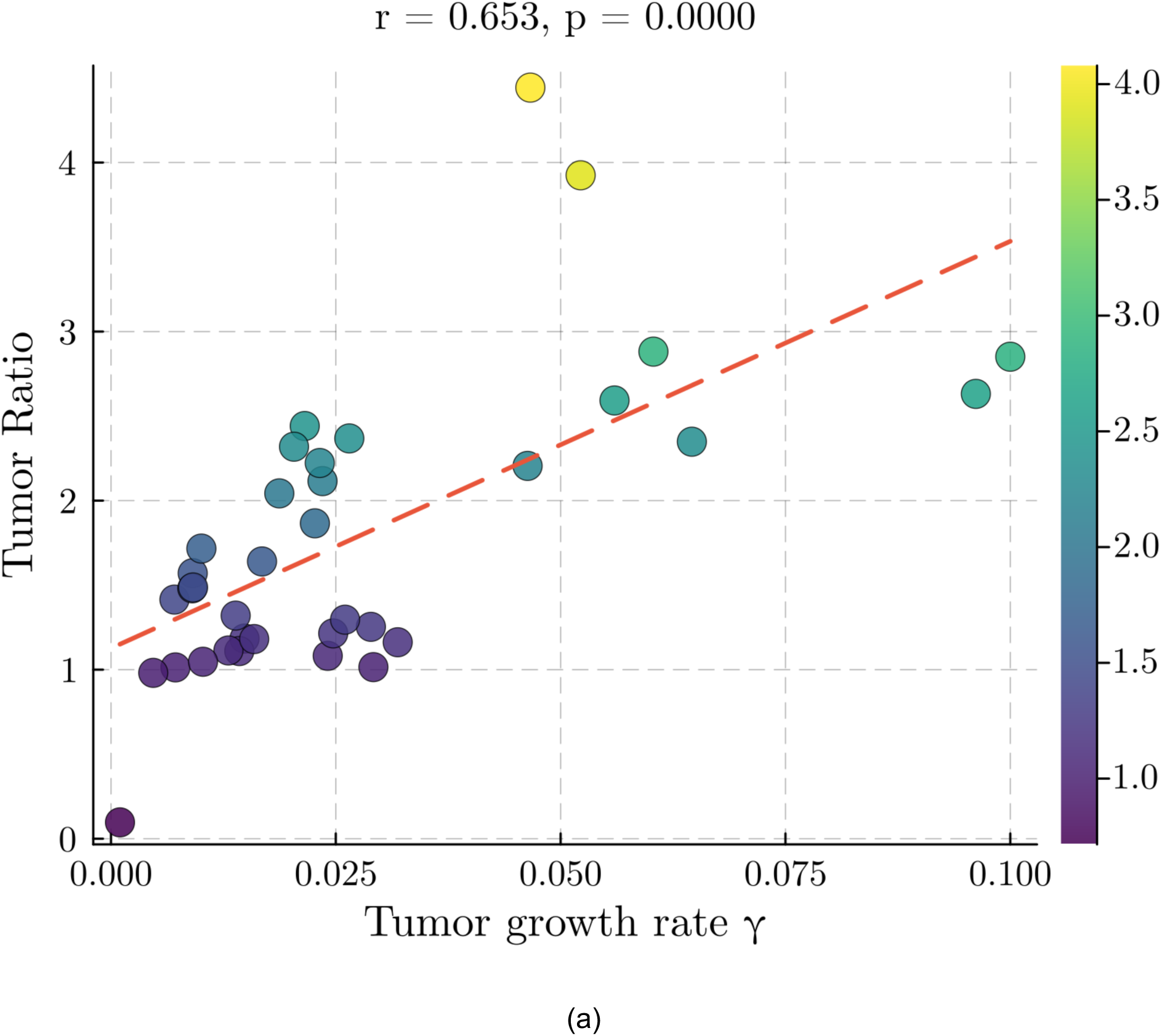
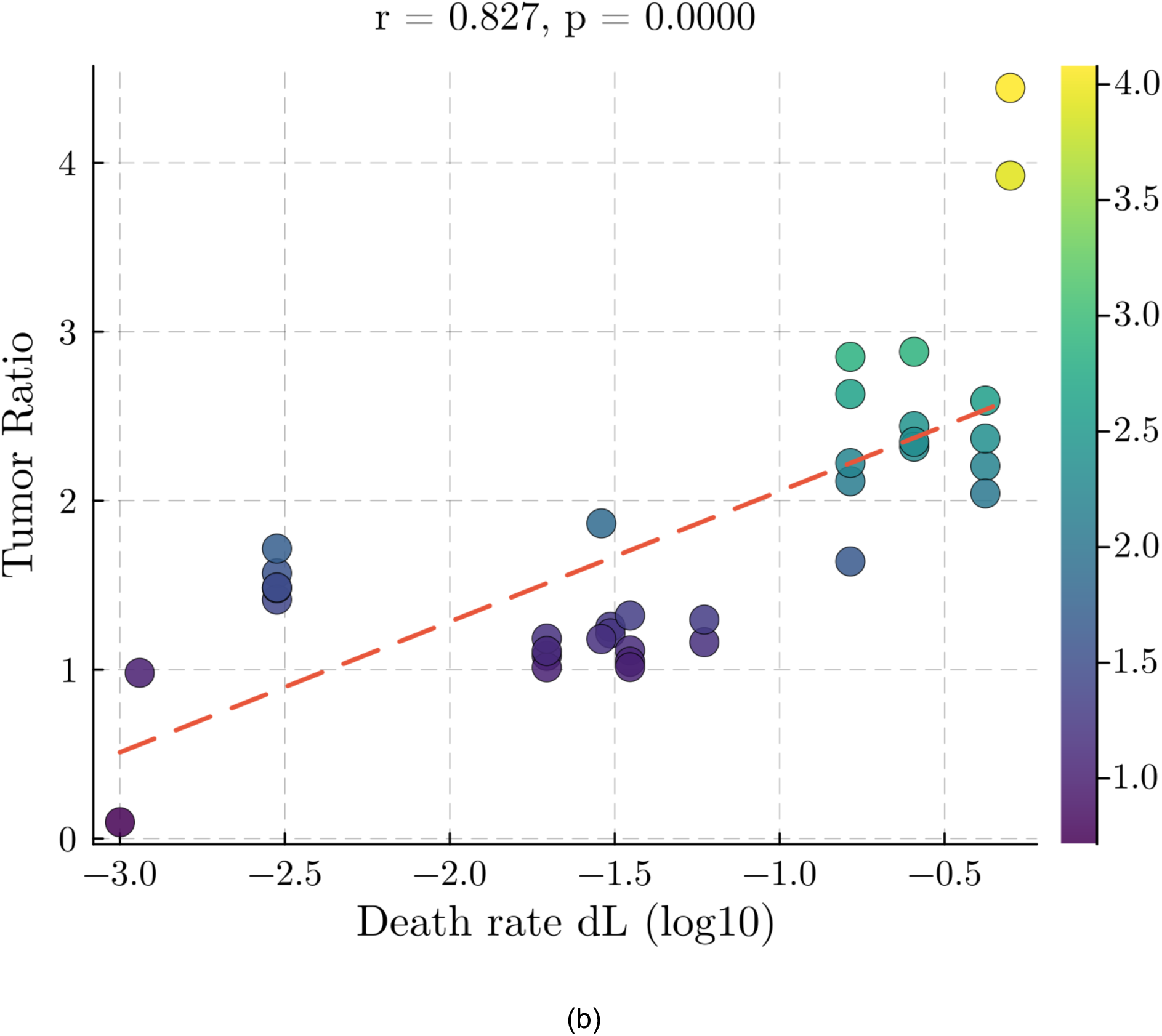
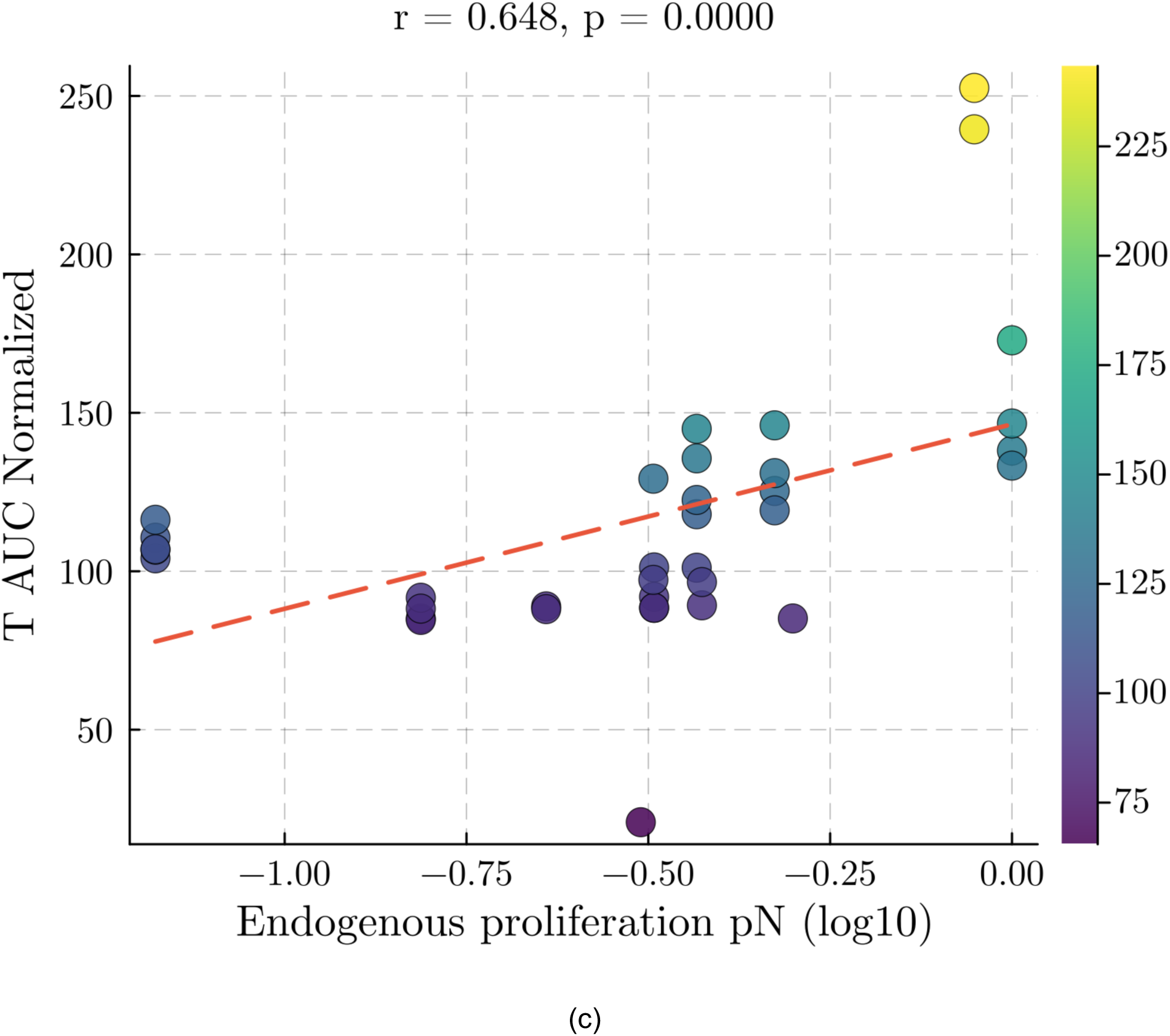
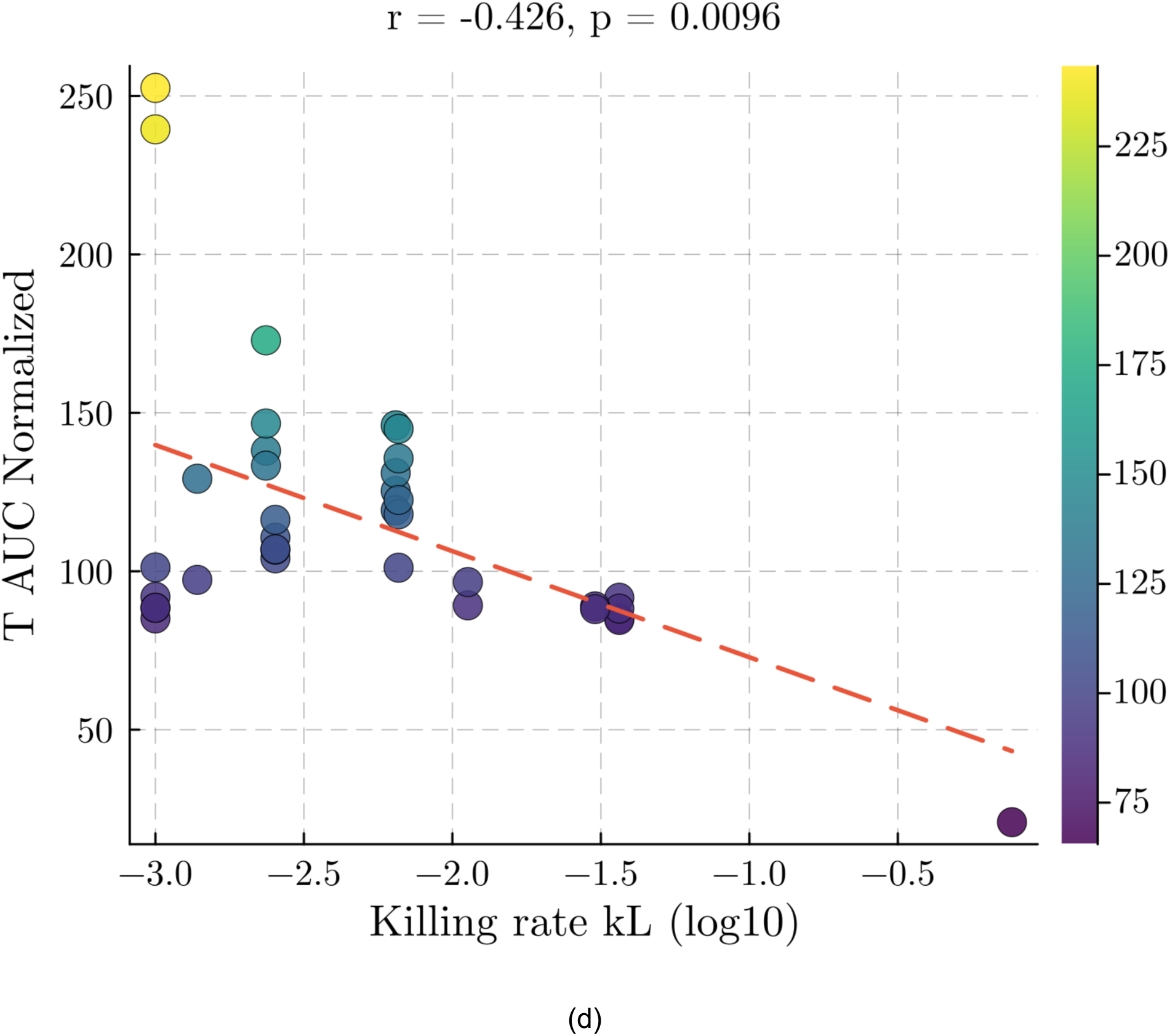
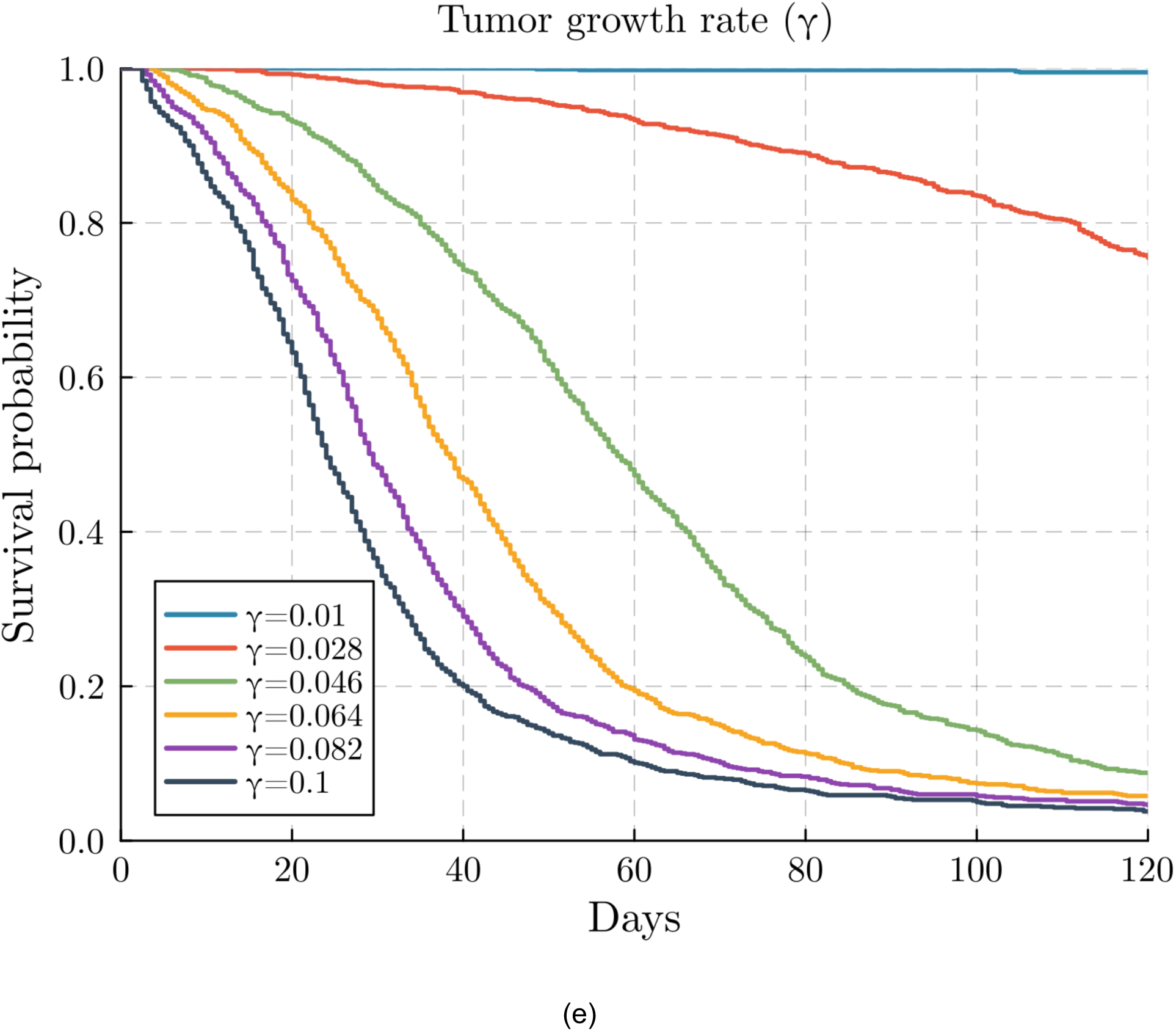
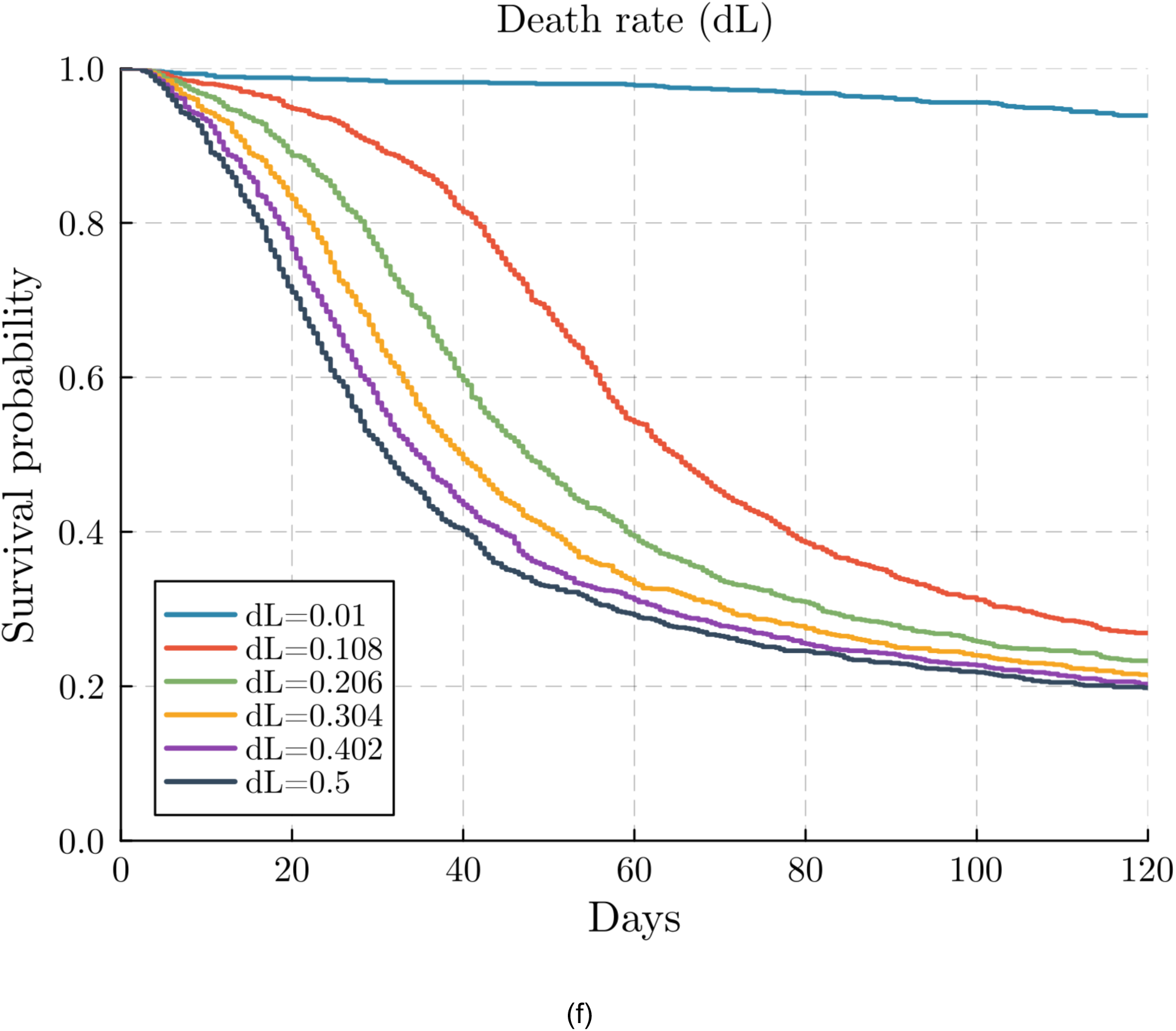
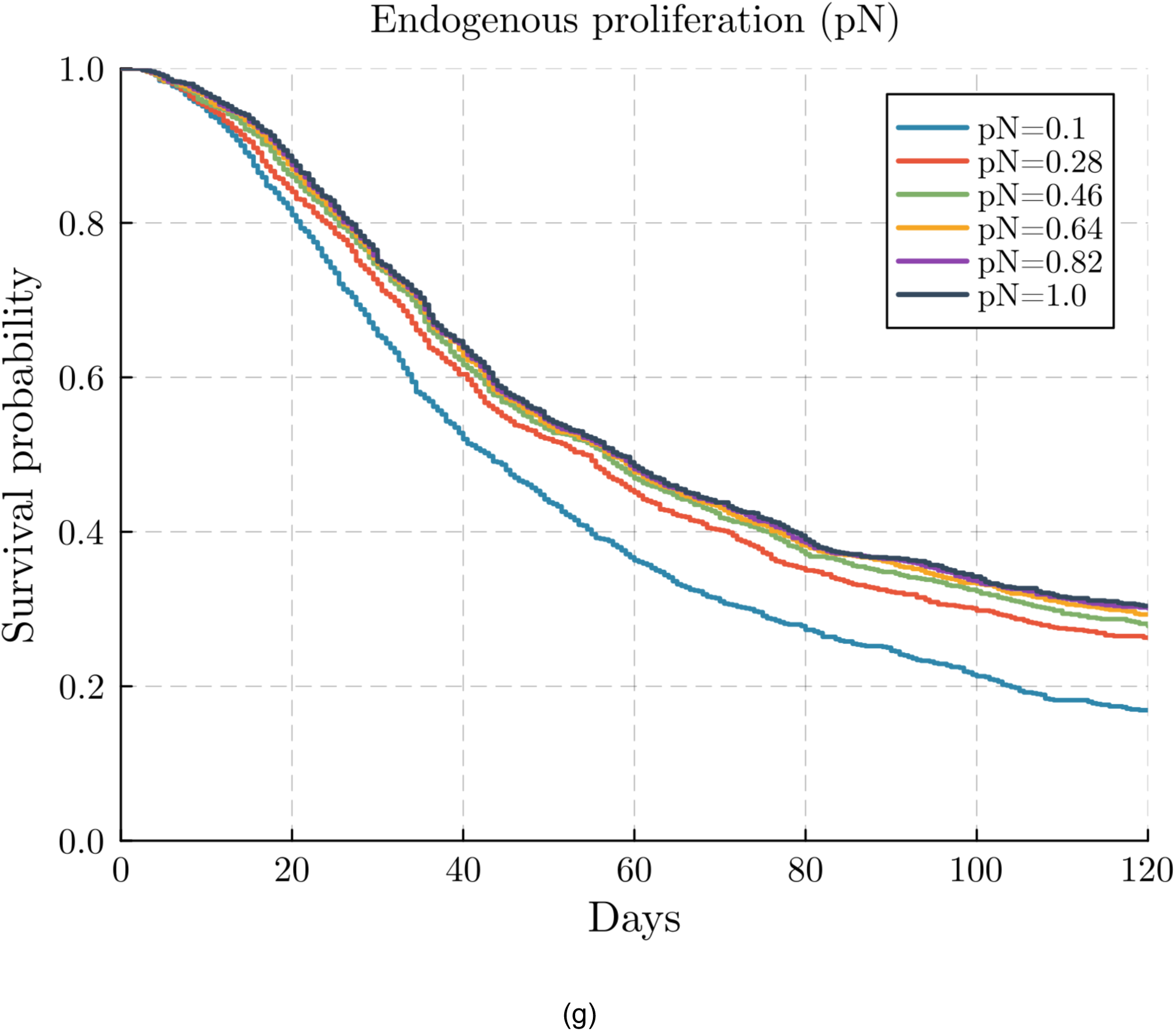
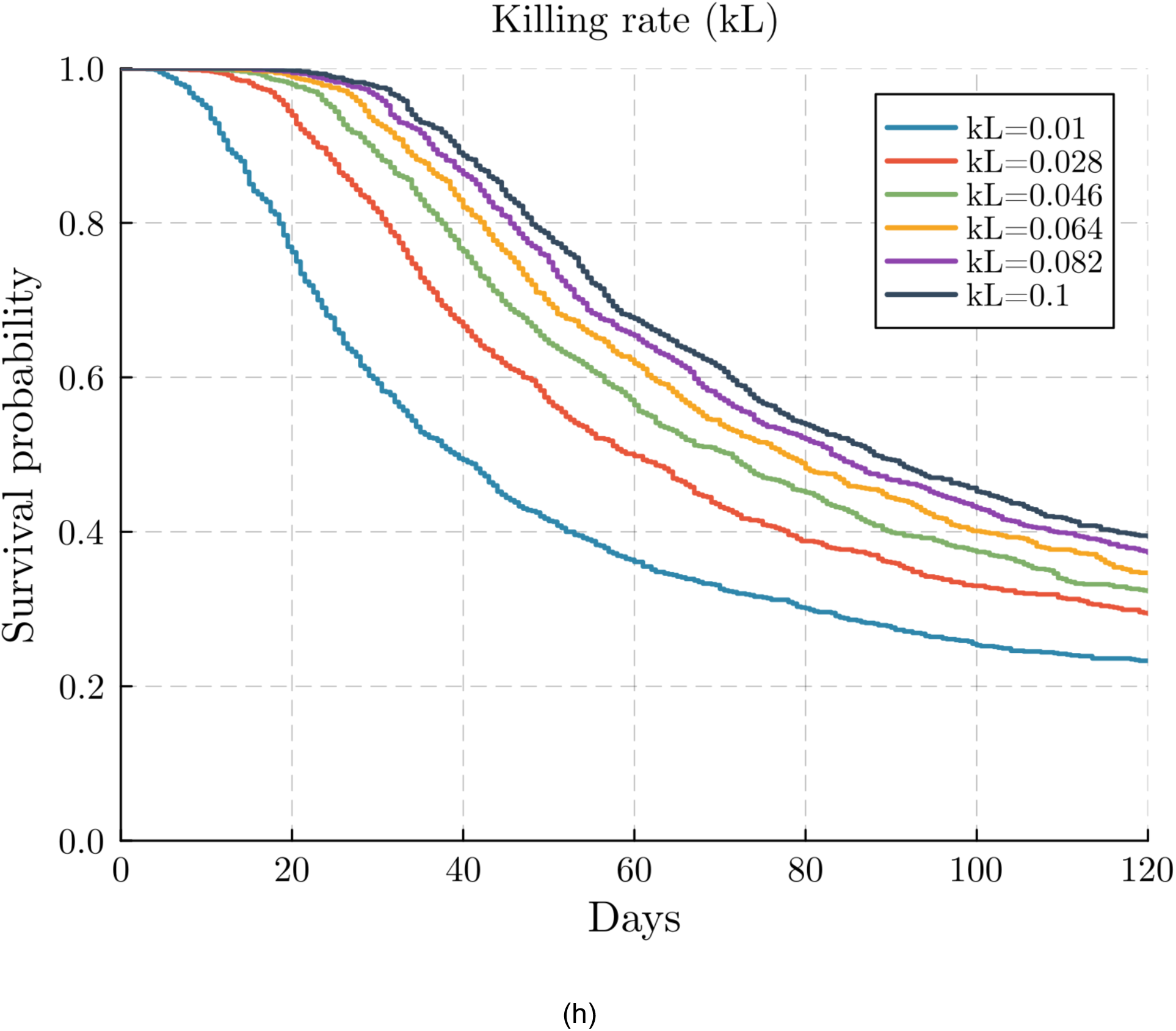
:**Tumor growth rate, T-cell persistence, endogenous T-cell competition, and cytotoxic potency are dominant determinants of therapeutic outcome. (a–d)** Scatter plots illustrating associations between fitted model parameters and therapeutic outcome metrics across all 27 tumor lesions in the clinical cohort. **(a)** Intrinsic tumor growth rate γ versus final tumor ratio (r = 0.653, p < 10^−4^). (b) *CISH-KO* T-cell death rate d_L_ versus final tumor ratio (r = 0.827, p < 10^−4^). **(c)** Endogenous T-cell proliferation rate p_N_ versus normalized tumor AUC (r = 0.648, p < 10^−4^), indicating that competitive expansion of endogenous T-cells is associated with worse therapeutic outcomes. (d) *CISH-KO* T-cell killing rate k_L_ versus normalized tumor AUC (r = −0.426, p = 0.0096). Points are colored by the corresponding secondary outcome metric (color bars at right); red dashed lines denote linear regression fits. **(e–h)** Kaplan–Meier survival curves from in silico experiments in which a single model parameter is varied while all others are held fixed at baseline values: **(e)** γ, **(f)** d_L_, **(g)** p_N_, and **(h)** k_L_. Each curve corresponds to a simulated cohort of 1000 patients; a death event is defined as tumor size exceeding 1.2× the initial size. Lower γ, lower d_L_, lower p_N_, and higher k_L_ each independently improve survival probability.

The associations between γ and k_L_ and therapeutic outcomes are mechanistically straightforward: faster-growing tumors impose a higher threshold for immune-mediated control, while greater per-cell cytotoxicity enables more effective tumor clearance. These two parameters define the fundamental balance between tumor expansion and immune suppression that determines whether therapy can achieve durable control.

In contrast, *CISH*-KO T-cell persistence (d_L_) emerged as the single strongest predictor of therapeutic failure. Biologically, rapid clearance of infused *CISH*-KO T-cells compresses therapeutic activity into a narrow temporal window, after which declining effector numbers cannot maintain sustained tumor control despite high per-cell cytotoxicity. This identifies *CISH*-KO T-cell persistence as a critical determinant of clinical response, which motivates the fractionated dosing strategy explored in the next section, where repeated infusions are simulated as a means to sustain effector cell numbers and extend the window of therapeutic activity.

Endogenous T-cell reconstitution (p_N_) emerged as the second-strongest predictor of therapeutic failure, revealing the unexpected consequence that faster host immune recovery paradoxically worsens treatment outcomes. This competitive displacement occurs because endogenous T-cells, despite substantially lower per-cell killing capacity than infused *CISH*-KO cells (k_N_ ≪ k_L_), rapidly reconstitute following lymphodepletion and occupy the shared immune niche. We note that p_N_ was not directly measured but inferred through model fitting; this association is a modeling-derived hypothesis requiring prospective experimental validation.The functional consequence is displacement of the engineered therapeutic population by less potent endogenous effectors.

This finding inverts the standard immunotherapy paradigm, which typically motivates supporting host immune reconstitution; here, aggressive endogenous T-cell recovery directly undermines engineered cell therapy efficacy.

To assess the causal implications of these associations, we performed in silico survival analyses using a simulated patient cohort generated by systematically varying each parameter while holding all others fixed (Figure 6e–h). These simulations confirm that reducing tumor aggressiveness (γ), enhancing *CISH*-KO T-cell persistence (lower d_L_), limiting endogenous T-cell competition (lower p_N_), and increasing cytotoxic potency (higher k_L_) each independently improve survival probability.

Collectively, these results indicate that clinical outcomes are governed by a balance between intrinsic tumor growth and multiple, mechanistically distinct properties of both the infused *CISH*-KO T-cells and the endogenous immune compartment. Notably, parameters controlling *CISH*-KO T-cell persistence (d_L_) and endogenous competition (p_N_) emerge as comparably important to cytotoxic strength (k_L_), suggesting that limited durability of infused *CISH*-KO T-cells and competitive displacement by less effective endogenous populations are major bottlenecks of current therapy. These findings point to complementary strategies for improving *CISH*-KO TIL efficacy: enhancing *CISH*-KO T-cell survival programs, augmenting killing efficiency, and mitigating competitive interference from endogenous T-cell reconstitution.

### Combination therapy improves tumor control without excessive T-cell exposure

The preceding analysis identified *CISH*-KO T-cell persistence (d_L_), endogenous T-cell competition (p_N_), and cytotoxic potency (k_L_) as key bottlenecks limiting the efficacy of *CISH*-KO TIL monotherapy. Motivated by murine studies showing that the combination of *CISH*-KO T-cells with anti–PD-1 checkpoint blockade restores complete tumor clearance where neither agent alone suffices, we used the calibrated model to explore whether combination strategies can address these bottlenecks in the clinical setting.

A mechanistic rationale for this combination arises from the biology of *CISH* disruption itself. *CISH* deficiency upregulates PD-1 expression on T-cells (14), rendering *CISH*-KO T-cells particularly susceptible to PD-1/PD-L1–mediated suppression within the tumor microenvironment. Anti–PD-1 blockade may therefore specifically benefit *CISH*-KO T-cells by relieving this checkpoint constraint, enhancing their functional efficiency—through sustained cytotoxicity and reduced exhaustion-driven attrition—rather than simply amplifying T-cell expansion. From the perspective of the bottlenecks identified above, this mechanism directly mitigates the persistence limitation by extending the effective lifespan of the therapeutic population.

We simulated six treatment scenarios: no therapy, standard *CISH*-KO TIL therapy, double-dose *CISH*-KO TIL therapy, anti–PD-1 monotherapy, *CISH*-KO TILs combined with anti–PD-1, and double-dose *CISH*-KO TILs combined with anti–PD-1. Model predictions are summarized in Figure 7(a) and (b).

**Figure 7.**
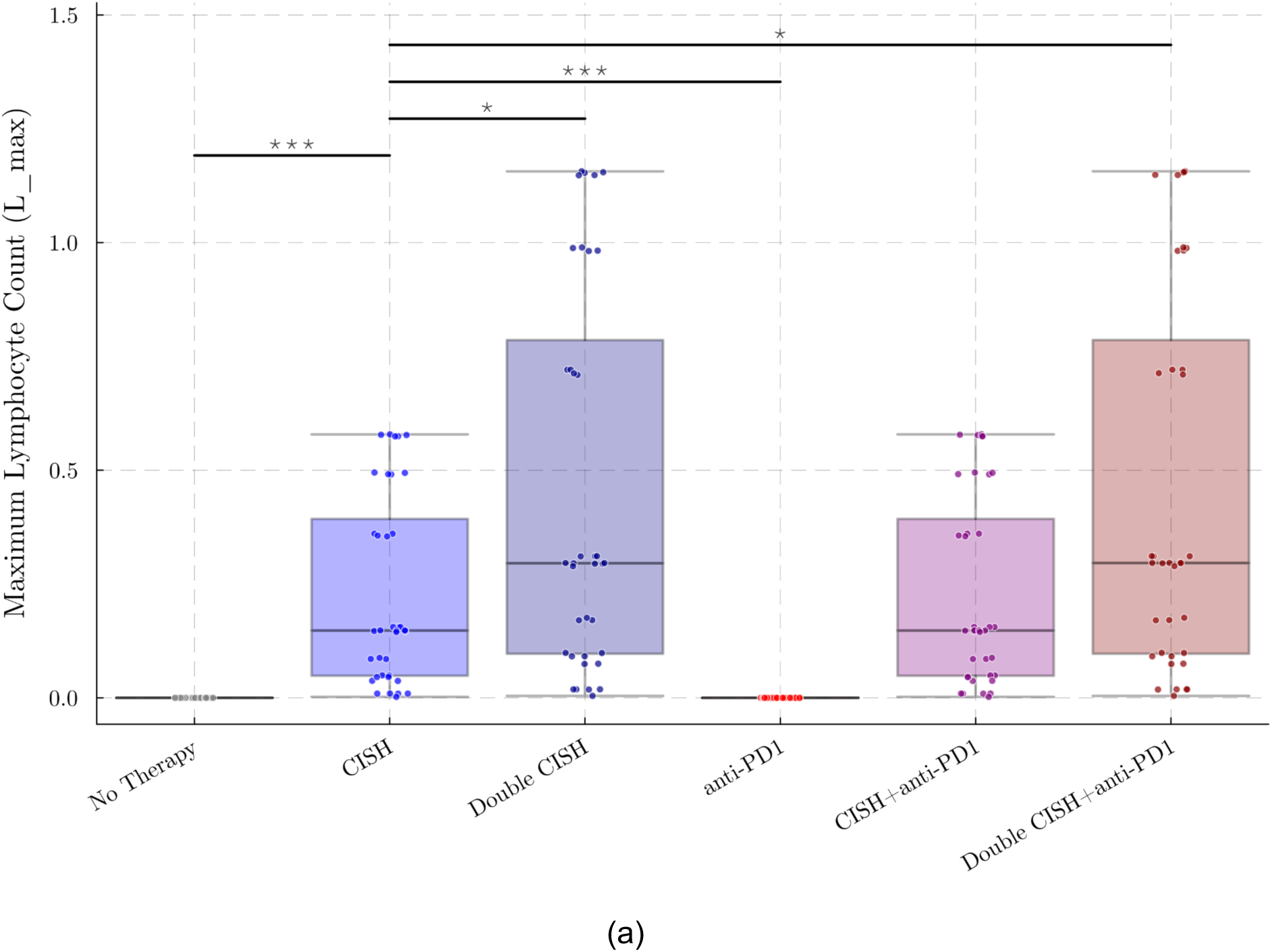
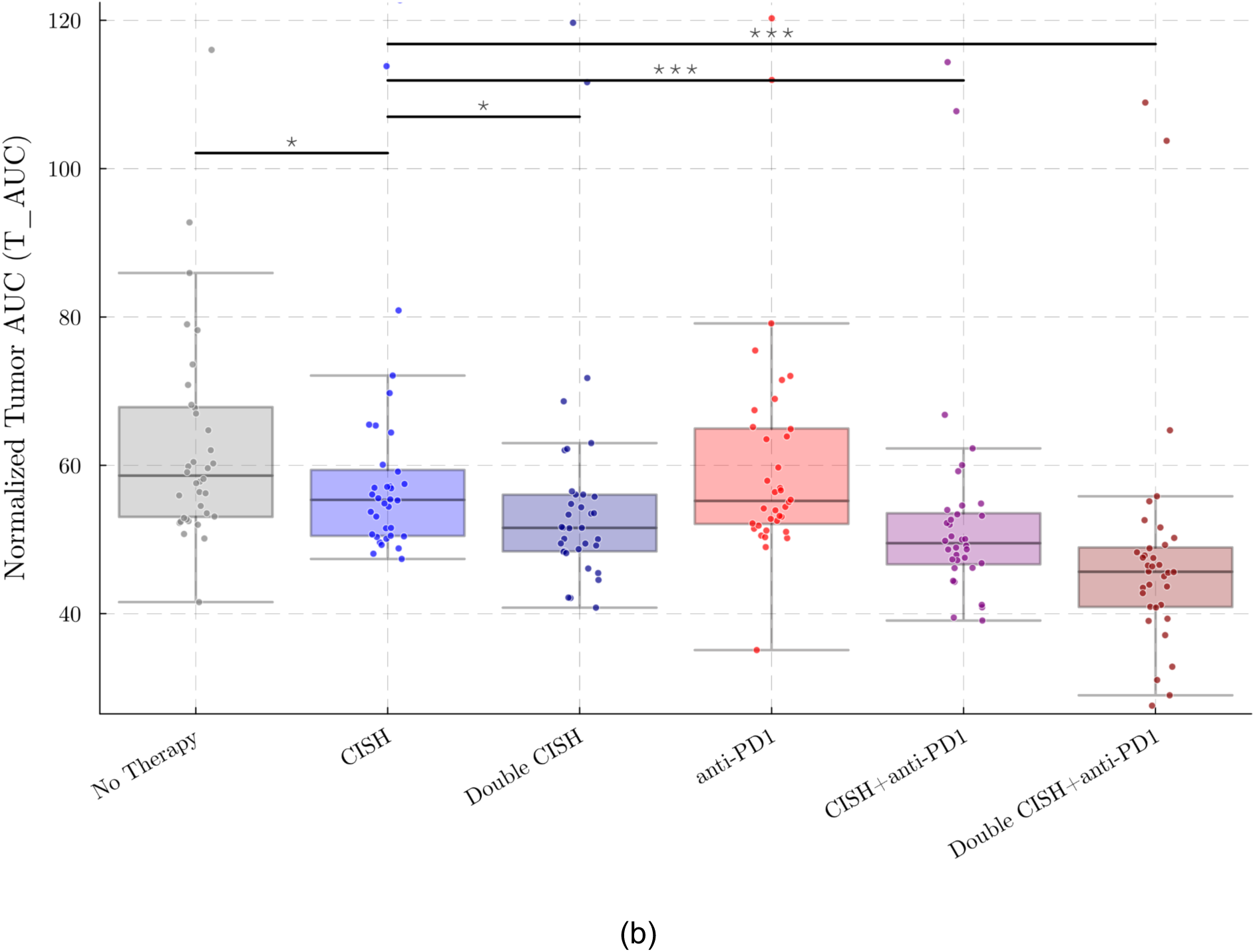
: Predicted therapeutic improvement under combination regimens. Model-predicted distributions of key therapeutic outcomes across six treatment scenarios: no therapy, *CISH-KO* TIL therapy, double-dose *CISH-KO* TIL therapy, anti–PD-1 monotherapy, *CISH-KO* TILs combined with anti–PD-1, and double-dose *CISH-KO* TILs combined with anti–PD-1. (a) Maximum effector T-cell burden (L_max_). (b) Normalized tumor exposure (T_AUC_). Combination therapies consistently outperform monotherapies, yielding both enhanced effector persistence and reduced tumor burden, consistent with synergistic mitigation of T-cell exhaustion and increased cytotoxic capacity.

Anti–PD-1 monotherapy alone produced minimal tumor suppression, consistent with experimental observations and with our finding that endogenous T-cells possess insufficient cytotoxic capacity (k_N_ ≪ k_L_) to control tumor growth regardless of checkpoint relief. Increasing the *CISH*-KO TIL dose two-fold substantially reduced tumor exposure but was accompanied by markedly elevated peak *CISH*-KO T-cell levels, raising potential concerns about toxicity without addressing the underlying persistence bottleneck. In contrast, the combination of standard-dose *CISH*-KO TILs with anti–PD-1 achieved comparable or superior tumor control while maintaining a lower maximum *CISH*-KO T-cell burden.

This result has two important implications. First, it demonstrates that improving the functional efficiency of existing *CISH*-KO T-cells is more effective than dose escalation alone, reinforcing the conclusion from our parameter analysis that persistence and per-cell potency—rather than sheer cell number—are the rate-limiting factors for therapeutic success. Second, the lower peak *CISH*-KO T-cell levels under combination therapy suggest a potentially more favorable safety profile, as excessive effector cell expansion has been associated with toxicity in adoptive cell therapy. These results support the rational design of combination regimens that enhance the durability and functional capacity of infused *CISH*-KO T-cells, rather than relying on dose intensification to overcome the biological barriers identified in this study.

### Fractionated dosing improves tumor control by prolonging *CISH*-KO T-cell persistence

The association between *CISH*-KO T-cell persistence and therapeutic outcome suggests that the temporal availability of infused *CISH*-KO T-cells may be an important determinant of response. A fractionated regimen, in which the total *CISH*-KO T-cell product is distributed across multiple infusions rather than delivered as a single bolus, could help maintain *CISH*-KO T-cells abundance over a longer interval. From the perspective of our model, this strategy may replenish the *CISH*-KO T-cell population before it falls below the level required for tumor control, thereby extending the duration of effective cytotoxic pressure without necessarily producing the high peak *CISH*-KO T-cell abundance associated with dose escalation.

Clinical studies on fractionated dosing have already been explored in the CAR-T setting. In particular, Frigault et al. (27) reports that fractionated CAR-T dosing can improve therapeutic outcomes in some settings while controlling peak cytokine levels and reducing the risk of severe cytokine release syndrome (CRS). Although CAR-T and *CISH*-KO TIL therapies differ biologically, these findings support the broader principle that dose scheduling can influence both efficacy and toxicity. Together with our model-based analysis, this motivates fractionated *CISH*-KO TIL delivery as a testable strategy for improving durable tumor control.

To test this hypothesis quantitatively, we used the calibrated patient-specific models to compare a single-bolus infusion against fractionated schedules that distribute the same total *CISH*-KO TIL dose across multiple infusions over a 56-day window. For each patient and schedule, we evaluated durable control via the end-to-start tumor size ratio and total tumor exposure via the cumulative AUC, expressing both as percent change from each patient’s single-bolus baseline so that heterogeneous responders could be compared on a common scale (Figure 8(a), (b)).

**Figure 8.**
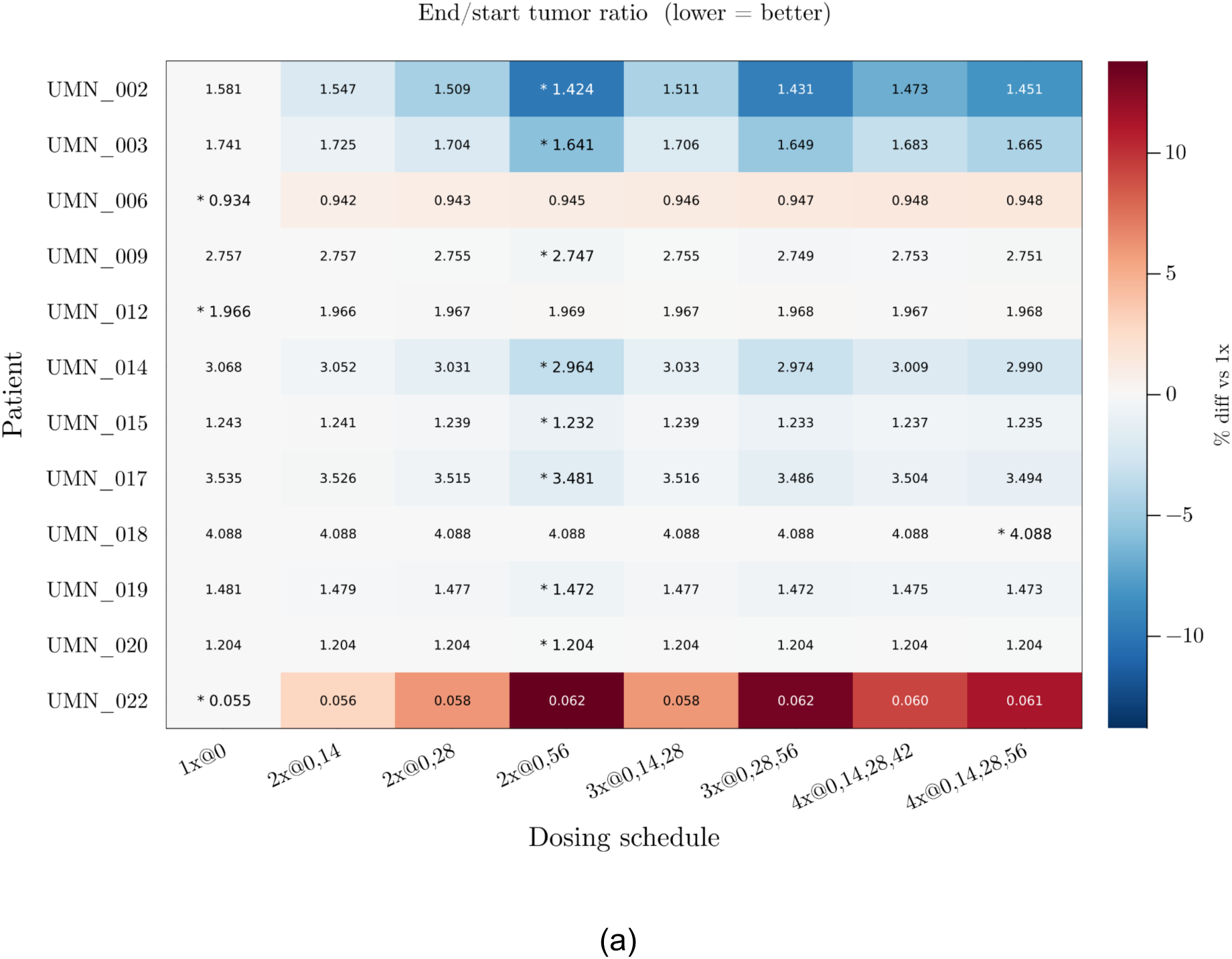
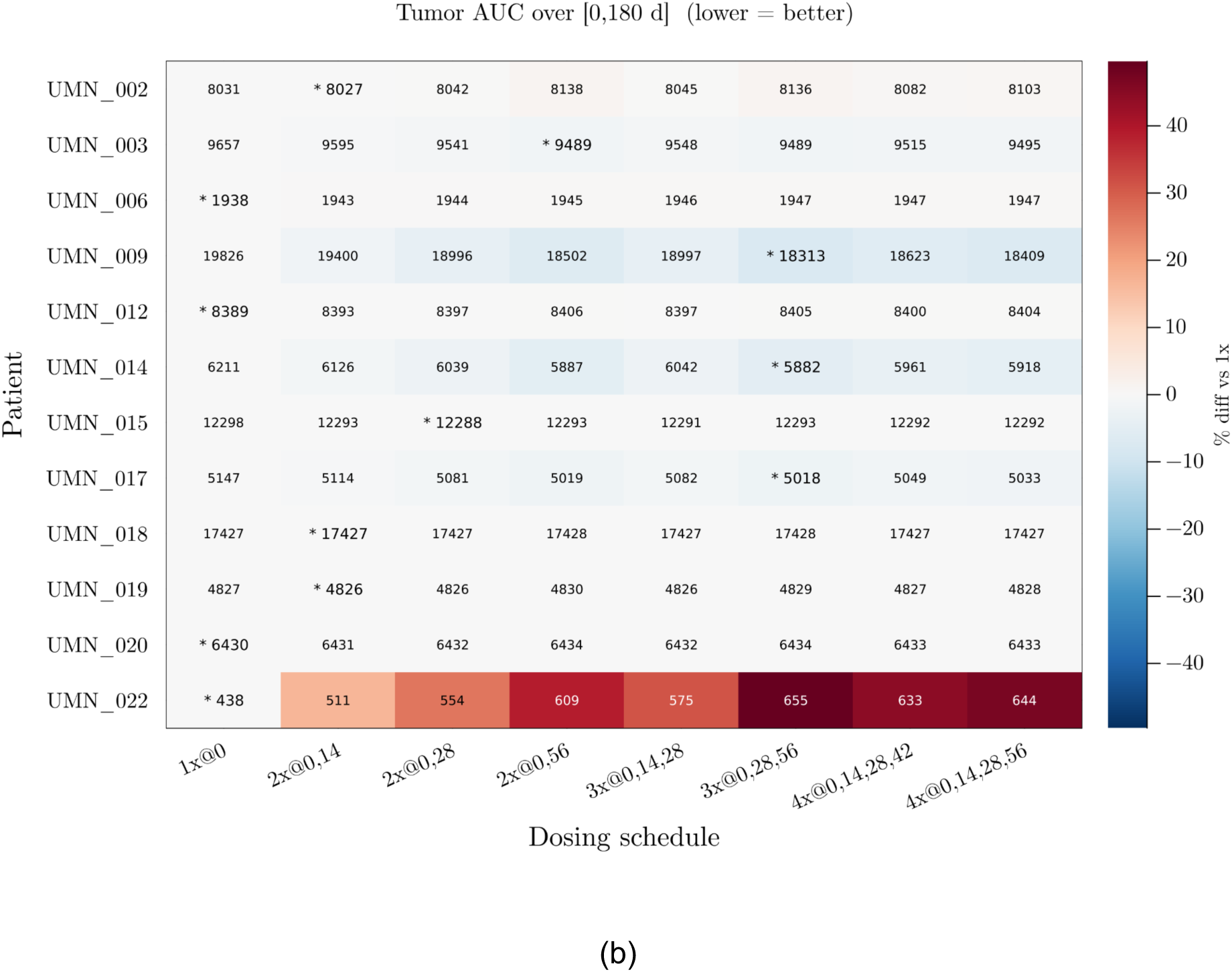
: **Predicted effect of dose fractionation on tumor control across the patient cohort**. For each of 12 patients, the calibrated patient-specific model was simulated over 180 days under eight dosing schedules sharing the same total CISH-KO TIL dose: a single bolus (1× @ 0) and seven fractionated regimens splitting the dose across 2–4 infusions spaced over a 56-day window. Cells are colored by percent change from each patient’s single-bolus baseline (blue = better, red = worse); absolute values are annotated and the row-wise best schedule is starred. (a) End-to-start tumor size ratio T(180)/T(0). (b) Cumulative tumor AUC over [0, 180 d].

The model predicts that fractionation produces consistent—if patient-dependent—improvements over single-bolus delivery. Two-dose regimens with the second infusion delayed to day 56, and three-dose regimens adding infusions at day 28 and day 56, most frequently emerge as the best schedule, while further subdivision into four fractions yields diminishing returns. The magnitude of benefit varies across patients but follows a clear pattern: patients whose fitted dynamics indicate that the tumor cannot be suppressed by a single bolus tend to show a meaningful benefit from fractionation, whereas patients already projected toward durable response or refractory progression are largely insensitive to schedule. UMN_006 and UMN_022 illustrate this insensitivity; in fact, for UMN_022 fractionation noticeably worsens the predicted outcome. Across the cohort, however, fractionation alone does not flip the qualitative therapeutic outcome for any patient—turning a non-responder into a responder appears to require combination therapy rather than dose scheduling.

Biologically, these results reinforce the persistence-limited picture of *CISH*-KO TIL therapy. A single bolus produces a transient effector peak that is rapidly attenuated by the per-cell decay rate, after which residual tumor regrows without effective cytotoxic pressure. Fractionation supplies fresh *CISH*-KO T-cells as the original infusion is being depleted, sustaining the cytotoxic-to-tumor ratio above the threshold for net tumor decline for a longer interval. Importantly, because fractionation redistributes rather than escalates the total dose, these gains are achieved without raising peak effector burden, offering a second lever—independent of combination therapy—for improving the therapeutic index of *CISH*-KO TIL therapy.

## Discussion

In this study, we developed a mechanistic mathematical model to characterize tumor response to *CISH*-KO TIL therapy across biological scales. By modeling interactions among tumor cells, endogenous TILs, and infused *CISH*-KO or wild-type T-cells, the framework recapitulated outcomes from two murine experiments and a first-in-human gastrointestinal cancer trial. Murine calibration revealed a conserved hierarchy of cytotoxic efficacy (k_L_ > k_Lo_ > k_N_), confirming the superior per-cell killing capacity of *CISH*-KO T-cells. The model was subsequently fit to longitudinal data from 27 tumors across 12 patients in a recent Phase 1 trial (24). Most patient tumors plateaued in size, representing stable disease, while the model predicts eventual tumor rebound for the majority of cases. The exception was patient UMN022, who achieved a durable complete clinical response that now exceeds three years since treatment; in this patient, the fitted killing rate k_L_ was the highest in the cohort by more than an order of magnitude, illustrating the quantitative threshold of cytotoxic potency required to overcome intrinsic tumor growth.

A central finding of the clinical analysis is that therapeutic outcomes are governed by four mechanistically distinct parameters: the intrinsic tumor growth rate (γ), the *CISH*-KO T-cell death rate (d_L_), the endogenous T-cell proliferation rate (p_N_), and the *CISH*-KO T-cell killing rate (k_L_). While γ and k_L_ define the intuitive balance between tumor expansion and immune suppression, the roles of d_L_ and p_N_ reveal less obvious but equally important barriers to durable response.

The strong association between d_L_ and treatment failure highlights *CISH*-KO T-cell persistence as a dominant bottleneck. This suggests that while *CISH*-KO enhances per-cell cytotoxic potency, its benefits are curtailed by the limited functional lifespan of the infused population(28,29). One possible contributor is the Rapid Expansion Protocol used in clinical manufacturing, which has been shown to preferentially expand clonotypes distinct from the tumor-reactive exhausted CD8+ population (30); the lower p_L_/p_N_ ratio observed in the clinical fits relative to mouse experiments would be consistent with such an effect, though our data cannot establish this link directly and other species-level differences may also contribute.Strategies to extend *CISH*-KO T-cell durability—such as optimizing *ex vivo* manufacturing to preserve a less differentiated phenotype (31), or co-administering cytokine support (32–34)—may therefore substantially improve clinical outcomes.

Equally notable is the finding that rapid endogenous T-cell reconstitution, captured by p_N_, is associated with worse therapeutic outcomes. This competitive displacement arises because endogenous T-cells, despite their markedly lower cytotoxic capacity (k_N_ ≪ k_L_), compete with the more potent *CISH*-KO population for limited resources within the tumor microenvironment, including cytokines, antigen presentation, and physical space. This modeling hypothesis requires prospective experimental validation, but is consistent with prior evidence from CAR-T therapy demonstrating that lymphodepletion depth and rate of endogenous immune recovery impacts treatment response and engineered T cell persistence (35,36). Together, this suggests that lymphodepletion timing and intensity could be further optimized to preserve niche availability for the therapeutic TIL population and enhance treatment efficacy.

Our in silico combination and fractionation strategies provide further support for addressing these bottlenecks through rational treatment design. Pairing *CISH*-KO TIL therapy with anti–PD-1 blockade achieved comparable or superior tumor control relative to dose escalation, while maintaining lower peak T-cell levels. This is mechanistically consistent with the biology of *Cish* disruption: because *Cish* deficiency upregulates PD-1 expression on T-cells, anti–PD-1 blockade specifically relieves checkpoint-mediated suppression of the therapeutic population, enhancing functional efficiency rather than simply increasing cell number. In contrast, anti–PD-1 monotherapy produced minimal benefit, consistent with our finding that endogenous T-cells lack sufficient cytotoxic capacity to control tumors regardless of checkpoint relief. These results reinforce that improving *CISH-KO* T-cell durability and efficiency is more impactful than dose escalation alone, and support the rational design of combination regimens tailored to the specific biological limitations of *CISH-KO* TIL therapy.

Fractionating the same total CISH-KO TIL dose across multiple delayed infusions offered a complementary, scheduling-based lever on the persistence bottleneck. The model predicted patient-dependent improvements over single-bolus delivery, with two- and three-dose regimens frequently emerging as the best schedule. It is notable that for one patient who did achieve a durable response to the treatment, dose fractionation worsened tumor ratio and tumor burden. Benefits were concentrated in patients whose single-bolus trajectory showed continued tumor progression, while those projected toward durable response or refractory progression were largely insensitive to schedule, and fractionation alone did not flip the qualitative outcome for any patient.

These gains are achieved without raising peak effector burden, providing a second lever—independent of combination therapy—for improving the therapeutic index of CISH-KO TIL therapy.

### Limitations

Several limitations should be considered when interpreting these findings. First, the ODE framework assumes well-mixed populations within the tumor microenvironment and does not capture spatial heterogeneity in T-cell infiltration, antigen distribution, or immunosuppressive gradients, all of which may influence local therapeutic efficacy. Second, the clinical cohort is small (n = 12 patients, 27 lesions), and the correlations in Figure 6a–d are computed across lesions nested within patients, introducing within-patient dependence that inflates the effective sample size. While the consistency of these associations across patients and their concordance with the in silico survival analyses (Figure 6e–h) provide supporting evidence, validation in larger, independent cohorts will be essential to confirm the generalizability of the identified parameter–outcome relationships and the four-parameter determinant structure. Third, circulating *CISH*-KO T-cell ratios were used as a proxy for intratumoral composition. Although peripheral and intratumoral T-cell frequencies have been shown to correlate in the early post-infusion period (25), differential trafficking, tissue retention, and local clonal dynamics may cause these compartments to diverge over time, particularly as circulating *CISH*-KO frequencies approach assay detection limits. Finally, the combination and fractionation predictions are derived entirely from in silico simulations calibrated to single-bolus monotherapy data and have not been validated in clinical or preclinical studies of these regimens. The anti–PD-1 enhancement parameters were estimated from murine data and transferred to the clinical model, assuming conserved PD-1 blockade pharmacodynamics across species. The simulated combination and scheduling outcomes should therefore be interpreted as qualitative predictions of the direction and relative magnitude of therapeutic benefit rather than quantitative forecasts of clinical response rates. Prospective clinical testing is needed to validate these predictions.

### Interpretation and clinical implications

Despite these limitations, the present analysis identifies actionable determinants of treatment response that can inform the design of next-generation TIL trials. While *CISH*-KO enhances cytotoxic potency, ultimate therapeutic success depends on whether *CISH*-KO T-cell persistence can outpace intrinsic tumor growth in the face of competitive pressure from the endogenous immune compartment. These findings suggest that strategies enhancing the survival, functional capacity, and competitive advantage of infused *CISH*-KO T-cells—such as simultaneous anti–PD-1 blockade, fractionated dosing of the same total cell product to extend effector persistence, optimized lymphodepletion regimens, and manufacturing protocols that preserve a less differentiated T-cell phenotype may be more effective than relying on dose escalation alone.

