## Supplement for "Modeling CISH-KO TIL therapy: T-cell persistence and endogenous competition determine response in gastrointestinal cancer"

### Supplementary Material

#### S1. Mouse Model

##### S1.0 Mathematical Model

As described in the main text, the model represents the tumor microenvironment through four interacting components: tumor cells ($T$), endogenous tumor-infiltrating lymphocytes (TILs; $N$), exogenously infused *CISH-KO* TILs ($L$), and exogenous wild-type TILs ($L_o$). Here we provide the full mathematical formulation, parameter definitions, and model selection rationale.

Tumor cells are assumed to proliferate exponentially at rate $\gamma$ and are eliminated through cytotoxic interactions with the three T-cell populations at rates $k_N$, $k_L$, and $k_{L_o}$, respectively. Each T-cell population undergoes natural death at rates $d_N$, $d_L$, and $d_{L_o}$, and tumor-induced exhaustion at rates $\mu_N$, $\mu_L$, and $\mu_{L_o}$.

T-cells proliferate at intrinsic rates $p_N$, $p_L$, and $p_{L_o}$. However, due to limited immune resources and spatial constraints (e.g., antigen availability and competition for niches), the three T-cell populations experience competitive suppression through a shared carrying capacity $K$. Logistic regulation of T-cell proliferation is a standard modeling framework for capturing resource-limited clonal expansion within confined tissue compartments, and has been widely employed in tumor–immune interaction models [1].

The assumption of a shared carrying capacity reflects competition among T-cell populations for common limiting resources within the tumor microenvironment, including cytokines (IL-2, IL-7, IL-15), antigen-presenting cell contacts, and spatial niches. Population-specific carrying capacities could in principle be introduced, but doing so would add parameters that are not identifiable from the available data. We therefore adopt the parsimonious single-$K$ formulation, noting that this implies symmetric resource competition among T-cell populations. Accordingly, T-cell proliferation is modulated by a logistic regulation term:

$$C(L, L_o, N) = 1 - \frac{L + L_o + N}{K}.$$

Endogenous TILs are additionally assumed to be recruited and expanded in response to tumor burden. This process is modeled via a tumor-stimulated accumulation term with rate $\delta$ and saturation level $h$, following the functional form introduced in [2].

**Anti–PD-1 therapy.** To model the effect of anti–PD-1 treatment on T-cell proliferation, cytotoxicity, and exhaustion inhibition [3], we introduce $Z$ to denote the concentration of anti–PD-1 drug. The pharmacokinetics of $Z$ are modeled as exponential decay with rate $\mu_Z$:

$$\frac{dZ}{dt} = -\mu_Z Z.$$

Following [4, 5], the enhancement of T-cell proliferation and the inhibition of tumor-induced exhaustion are modeled by multiplying or dividing the corresponding baseline terms by a factor $1 + Z$. In addition, motivated by the sigmoidal dependence of T-cell activity on PD-1/PD-L1 blockade [5], we model enhanced cytotoxicity using sigmoid functions $\alpha(Z)$ and $\alpha_L(Z)$ for wild-type and *CISH-KO* T-cells, respectively. This distinction reflects experimental evidence that *Cish* deficiency alters activation marker expression and susceptibility to PD-1 blockade [6]. Specifically,

$$\begin{aligned}\alpha(Z) &= \frac{a+1}{a+e^{-bZ}},\\ \alpha_L(Z) &= \frac{a+1}{a+e^{-b_L Z}},\end{aligned}$$

where $a$, $b$, and $b_L$ control the shape of the sigmoidal response. Both $\alpha(Z)$ and $\alpha_L(Z)$ are monotone increasing and satisfy $\alpha(0) = \alpha_L(0) = 1$.

**Adoptive T-cell dosing.** To model the pharmacokinetics of exogenously infused T-cells, we link the administered dose to the effector populations $L$ and $L_o$ through an intermediate dose compartment. Specifically, the administered dose decays according to

$$\begin{aligned}\frac{d\text{Dose}}{dt} &= -f_{\text{dose}}\,\text{Dose} - \kappa\,\text{Dose},\\ \frac{dL_I}{dt} &= \kappa\,\text{Dose},\\ \frac{d\text{Dose}_o}{dt} &= -f_{\text{dose}}\,\text{Dose}_o - \kappa\,\text{Dose}_o,\\ \frac{dL_{o,I}}{dt} &= \kappa\,\text{Dose}_o,\end{aligned} \tag{1}$$

where $L_I$ and $L_{o,I}$ respectively represent the *CISH-KO* T-cells and wild-type T-cells infused from outside. Here, $f_{\text{dose}}$ represents loss of administered cells due to inefficient engraftment or clearance prior to tumor infiltration, while $\kappa$ controls the rate at which infused T-cells successfully infiltrate the tumor microenvironment and contribute to the effector TIL populations.

**Full mouse model.** Under the simplification $\mu_{L_o} = \mu_N$, $d_{L_o} = d_N$, and $p_{L_o} = p_N$ (see Section S1.2), the mouse experiments employ a seven-compartment ODE system:

$$\begin{aligned}
\frac{d\,\mathrm{Dose}_L}{dt} &= -\kappa\,\mathrm{Dose}_L, \\
\frac{d\,\mathrm{Dose}_{L_o}}{dt} &= -\kappa\,\mathrm{Dose}_{L_o}, \\
\frac{dT}{dt} &= \gamma\,T - \alpha_L(Z)\,k_L\,L\,T - \alpha(Z)\,k_{L_o}\,L_o\,T - \alpha(Z)\,k_N\,N\,T, \\
\frac{dL}{dt} &= -\frac{\mu_L\,L\,T}{1+Z} - d_L\,L + C(L, L_o, N)\,p_L\,L\,(1+Z) + \kappa\,\mathrm{Dose}_L, \\
\frac{dL_o}{dt} &= -\frac{\mu_N\,L_o\,T}{1+Z} - d_N\,L_o + C(L, L_o, N)\,p_N\,L_o\,(1+Z) + \kappa\,\mathrm{Dose}_{L_o}, \\
\frac{dN}{dt} &= -\frac{\mu_N\,N\,T}{1+Z} - d_N\,N + C(L, L_o, N)\,p_N\,N\,(1+Z) + C(L, L_o, N)\,\frac{\delta\,T}{h+T}, \\
\frac{dZ}{dt} &= -\mu_Z\,Z,
\end{aligned} \tag{2}$$

where $C(L, L_o, N) = 1-(L+L_o+N)/K$ is the logistic capacity term limiting total T-cell expansion, and the anti-PD-1 enhancement functions are

$$\alpha(Z) = \frac{a+1}{a+e^{-b\,Z}}, \qquad \alpha_L(Z) = \frac{a+1}{a+e^{-b_L\,Z}}. \tag{3}$$

The functions $\alpha(Z)$ and $\alpha_L(Z)$ modulate tumor killing efficacy in the presence of anti-PD-1 therapy. When $Z = 0$ (no anti-PD-1), both functions evaluate to 1 and the killing rates revert to their baseline values. As $Z$ increases, $\alpha$ and $\alpha_L$ increase monotonically, representing checkpoint-blockade-mediated enhancement of cytotoxicity. The separate steepness parameters $b$ and $b_L$ allow *CISH-KO* and wild-type/endogenous T-cells to respond differently to anti-PD-1 treatment. Similarly, the factor $(1+Z)$ in the exhaustion and proliferation terms captures the modulatory effect of anti-PD-1: exhaustion is attenuated by the factor $1/(1+Z)$, while proliferation is enhanced by the factor $(1+Z)$.

The patient model of Section S2 (Equation (6)) is obtained from (2) by removing the wild-type infused T-cell compartment ($L_o$), the drug compartment ($Z$), and adding a pre-engraftment clearance term $f_{\mathrm{dose}}\,\mathrm{Dose}$ to the dose equation.

#### S1.1 Data Sources and Experimental Design

We first calibrated the model using data from mouse experiments reporting longitudinal tumor size measurements under different therapeutic conditions. In the 2015 study [7], mice bearing established B16 melanoma tumors (tumor area $T_0 = 40$ mm$^2$ at day 0) were assigned to three groups: untreated controls (NT), mice receiving adoptively transferred wild-type T-cells (*Cish*$^+$), and mice receiving *CISH-KO* T-cells (*Cish*$^-$). Tumor area (mm$^2$) was measured longitudinally over a 32-day observation window.

The 2022 study [6] extended this design to six groups by further stratifying mice based on the presence or absence of anti-PD-1 treatment. Mice bearing established tumors ($T_0 = 25$ mm$^2$ at day 0) were randomized into: (1) no T-cells, isotype control; (2) no T-cells + anti-PD-1; (3) wild-type T-cells, isotype control; (4) wild-type T-cells + anti-PD-1; (5) *CISH-KO* T-cells, isotype control; (6) *CISH-KO* T-cells + anti-PD-1. Tumor area was measured longitudinally over a 42-day observation window. The inclusion of anti-PD-1 arms in the 2022 study activates the drug compartment $Z$ and the enhancement functions $\alpha(Z)$, $\alpha_L(Z)$ in (2), making the anti-PD-1-related parameters ($a$, $b$, $b_L$, $\mu_Z$) identifiable.

#### S1.2 Model Assumptions and Parameter Selection

**Initial conditions.** Because the administered T-cell dose was not reported in these experiments, we fixed the infused dose to 0.5 mm$^2$ and the initial anti-PD-1 concentration to $Z(0) = 5$ (dimensionless units) across all mouse simulations. As mice underwent lymphodepletion prior to adoptive cell transfer, all T-cell populations were initialized at zero at the time of infusion. All treatment groups within each study were assumed to share a common set of model parameters, with therapeutic effects captured exclusively through differences in treatment inputs rather than parameter values. The initial conditions for the 2015 and 2022 experiments are summarized in Table 1.

| **Treatment arm** | $T(0)$ | $L(0)$ | $L_o(0)$ | $N(0)$ | $Z(0)$ | $\text{Dose}_L(0)$ | $\text{Dose}_{L_o}(0)$ |
|---|---|---|---|---|---|---|---|
| *2015 study* ($T_0 = 40$ mm$^2$) | | | | | | | |
| NT | 40 | 0 | 0 | 0 | 0 | 0 | 0 |
| WT | 40 | 0 | 0 | 0 | 0 | 0 | 0.5 |
| *CISH-KO* | 40 | 0 | 0 | 0 | 0 | 0.5 | 0 |
| *2022 study* ($T_0 = 25$ mm$^2$) | | | | | | | |
| No cells | 25 | 0 | 0 | 0 | 0 | 0 | 0 |
| No cells + aPD1 | 25 | 0 | 0 | 0 | 5 | 0 | 0 |
| WT | 25 | 0 | 0 | 0 | 0 | 0 | 0.5 |
| WT + aPD1 | 25 | 0 | 0 | 0 | 5 | 0 | 0.5 |
| *CISH-KO* | 25 | 0 | 0 | 0 | 0 | 0.5 | 0 |
| *CISH-KO* + aPD1 | 25 | 0 | 0 | 0 | 5 | 0.5 | 0 |

Table 1: Initial conditions for each treatment arm in the 2015 and 2022 mouse experiments.

**Fixed parameters.** Baseline values for endogenous T-cell parameters were drawn from prior tumor–immune modeling studies [2, 8–10] and are listed in Table 2. In contrast, kinetic parameters governing engineered T-cell populations have been less extensively characterized in the literature and are therefore estimated from the experimental data in this work. Because the infused dose was fixed exogenously rather than derived from reported data, we did not model additional dose loss prior to tumor infiltration and set $f_{\text{dose}} = 0$. To further reduce model complexity and facilitate parameter identifiability, we assumed that wild-type engineered T cells share the same immune-related kinetic parameters as endogenous T cells, including exhaustion, natural decay, and proliferation rates (i.e., $\mu_{L_o} = \mu_N$, $d_{L_o} = d_N$, and $p_{L_o} = p_N$). The sole exception was the tumor killing rate, which was allowed to differ to reflect the enhanced tumor recognition conferred by the engineered T-cell receptor. The immune carrying capacity $K$ was set to 1 mm$^2$, corresponding to an overall tumor size on the order of 10–100 mm$^2$. The infiltration rate $\kappa$ was manually tuned for each study based on the observed timing of the transient tumor expansion prior to immune-mediated suppression: the earlier tumor peak in the 2015 study (approximately Day 5) required a larger $\kappa$, reflecting rapid T-cell accumulation, whereas the delayed peak in the 2022 data (approximately Day 8) was captured by a smaller $\kappa$, consistent with slower infiltration dynamics.

**Fitted parameters.** Given the constraints above, the parameters to be estimated are primarily those governing *CISH-KO* T-cell dynamics. In the 2022 study, the inclusion of anti–PD-1 treatment arms additionally makes the drug-related parameters identifiable.

In the 2022 study, nine parameters were estimated:

$$\Theta_{2022} = \{\gamma,\, k_L,\, k_{L_o},\, k_N,\, p_L,\, d_L,\, a,\, b,\, b_L\}.$$

In the 2015 study, four parameters were estimated:

$$\Theta_{2015} = \{\gamma,\, k_L,\, p_L,\, d_L\}.$$

The wild-type killing rate $k_{L_o}$ and endogenous T-cell killing rate $k_N$ estimated from the 2015 study were carried forward to the 2022 analysis as reference parameters to maintain consistency across the two datasets, reflecting the biological expectation that these killing rates should be comparable when employing the same T-cell products and tumor model.

Since no anti-PD-1 drug was administered in the 2015 study, the drug-related parameters ($a$, $b$, $b_L$, $\mu_Z$) were not identifiable and were neglected.

##### S1.3 Fitting Procedure

Model calibration was performed by minimizing a weighted sum of scaled squared errors between simulated and observed tumor sizes. For each dataset $d$ with observed tumor data $\{(t_k,\, T_k^{\text{obs}})\}_{k=1}^{K_d}$, the scaled squared error was computed as:

$$\mathcal{L}_d = \sum_{k=1}^{K_d} \left( \frac{T_d^{\text{obs}}(t_k) - T_d^{\text{model}}(t_k)}{S_d} \right)^2, \tag{4}$$

where $S_d = \max(\max(T_d^{\text{obs}}), 10)$ is the scale factor for each arm to normalize residuals relative to the observed maximum tumor size. This scale-invariant formulation ensures that arms with naturally smaller tumor burdens (e.g., successful treatment arms) contribute comparably to the objective function.

For both the 2015 and 2022 studies, the total objective is:

$$\mathcal{L}_{\text{total}} = \sum_{d} w_d \, \mathcal{L}_d, \tag{5}$$

where $w_d$ is the weight assigned to each treatment arm. For the 2015 study, weights are $(w_{\text{NT}}, w_{\text{WT}}, w_{\text{CISH KO}}) = (1.0, 2.0, 5.0)$. For the 2022 study, weights follow the ordering of treatment arms in Section S1.1 as $(w_1, w_2, w_3, w_4, w_5, w_6) = (1.0, 1.0, 2.0, 3.0, 5.0, 5.0)$, with larger weights assigned to arms containing infused T-cells (both WT and CISH-KO) to ensure balanced contribution across all conditions.

The ODE system was solved using the Tsit5 (Tsitouras 5th-order Runge–Kutta) method implemented in `DifferentialEquations.jl`, with absolute tolerance $10^{-10}$ and relative tolerance $10^{-3}$. Parameter optimization was carried out using the L-BFGS-B algorithm with box constraints, implemented in `Optim.jl`. A multistart strategy with $N_{\text{start}} = 20$ random initial points drawn uniformly from the feasible region $[\theta_{\text{lb}}, \theta_{\text{ub}}]$ was employed. All starts were executed in parallel using Julia's multi-threading facility, and the parameter set yielding the lowest objective value was selected. All optimization was implemented in Julia (v1.10+) using `DifferentialEquations.jl` for ODE integration and `Optim.jl` for optimization.

#### S2. Patient Model

In the clinical trial setting there are no wild-type infused T-cells and no anti-PD-1 drug; the general model therefore reduces to the following four-compartment system:

$$\begin{aligned}
\frac{d\,\text{Dose}}{dt} &= -f_{\text{dose}}\,\text{Dose} - \kappa\,\text{Dose}, \\
\frac{dT}{dt} &= \gamma\,T - k_L\,L\,T - k_N\,N\,T, \\
\frac{dL}{dt} &= -\mu_L\,L\,T - d_L\,L + C(L,N)\,p_L\,L + \kappa\,\text{Dose}, \\
\frac{dN}{dt} &= -\mu_N\,N\,T - d_N\,N + C(L,N)\,p_N\,N + C(L,N)\,\frac{\delta\,T}{h+T},
\end{aligned} \tag{6}$$

where $C(L, N) = 1 - (L + N)/K$ is the logistic capacity term limiting total T-cell expansion.

##### S2.1 Data Sources and Notation

Three patient-level data streams were used to calibrate the model. We index patients by $i = 1, \ldots, N_p$, tumor lesions within patient $i$ by $j = 1, \ldots, J_i$, and serial observation time points by $k$.

- **Tumor size data** $\{(t_{ik}, T_{ijk})\}$. Longitudinal tumor burden was measured via imaging for each patient $i$ at time points $t_{ik}$ (days post-infusion). The measurement $T_{ijk}$ denotes the observed tumor area ($\text{mm}^2$) of lesion $j$ at time $t_{ik}$. Patients harbored between 1 and 4 individually tracked lesions. The first observation for each lesion, $T_{ij1}$, was used as the tumor-specific initial condition $T_j(0) = T_{ij1}$ in the ODE system. These data directly inform the tumor component of the objective function, $\mathcal{L}_{\text{tumor}}$ (Equation (8)).

- ***CISH-KO*/wild-type T-cell ratio data** $\{(s_{ik}, R_{ik})\}$. The ratio of *CISH-KO* to wild-type T-cells was quantified by flow cytometry from peripheral blood samples collected at time points $s_{ik}$ (days post-infusion). The measurement $R_{ik}$ represents the observed ratio $L^{\text{obs}}/N^{\text{obs}}$ in circulation for patient $i$ at time $s_{ik}$. Only post-infusion samples ($s_{ik} \geq 0$) were included in model fitting. Because direct measurement of the *CISH-KO*/wild-type ratio among tumor-infiltrating lymphocytes is not feasible in vivo, we assume that the ratio within the tumor microenvironment mirrors the ratio observed in peripheral blood. This approximation is motivated by studies demonstrating that the persistence and expansion of adoptively transferred T-cell clonotypes in peripheral blood correlate with clinical tumor regression [11,12], suggesting that circulating T-cell dynamics are informative about therapeutic activity at the tumor site. The assumption may break down at later time points due to differential trafficking, tissue retention, and local clonal expansion; the fitting procedure accounts for this through reduced weighting and first-time-point normalization of the ratio component (see Section S2.3 for details). Under this assumption, the model-predicted ratio $\hat{R}(t) = \bar{L}(t)/\bar{N}(t)$ (averaged across tumors) can be directly compared to the blood-derived measurement $R_{ik}$, informing $\mathcal{L}_{\text{CISH}}$ (Equation (9)).

- **Infused T-cell dose** $D_i(0)$. Each patient received a single infusion of *CISH-KO* T-cells at day 0. The total number of infused cells, $N_{\text{infused}}$, was recorded per patient.

  The infused T-cell dose is typically on the order of $10^{10}$ cells, while the tumor burden variables in the model ($T$, $L$, and $N$) are represented in units of tumor area (mm$^2$). To keep the magnitudes of all state variables comparable within the ODE system, we rescale the infused cell number by a reference dose of $10^{10}$ cells, mapping it into the same tumor-area units (mm$^2$) used for the other state variables. Specifically, the dose compartment is initialized as

$$D_i(0) = \frac{N_{\text{infused}}}{10^{10}} \quad (\text{mm}^2).$$

This rescaling is absorbed into the infiltration rate $\kappa$ and pre-engraftment clearance rate $f_{\text{dose}}$, which jointly calibrate the effective rate and efficiency of T-cell delivery to each tumor site. Because the clinical data informing T-cell dynamics consist of the ratio between *CISH-KO* and endogenous T-cells rather than absolute cell counts, the choice of reference dose does not affect fitting performance or the results of the statistical analysis—the relative ordering and magnitude relationships among fitted parameters are preserved. The scaling does limit the direct biological interpretation of absolute T-cell state variable values, but this is not the focus of the present analysis. Similar rescaling approaches are standard in pharmacokinetic–pharmacodynamic modeling of adoptive cell therapies [8,9]. Biologically, only a small fraction of infused T-cells ultimately traffics to a given tumor lesion, while the majority remain in circulation or distribute to other tissues. Representing the circulating dose as a scaled quantity therefore serves as a practical approximation for modeling the pool of T-cells available for tumor infiltration.

#### S2.2 Parameter Selection and Initial States

**Initial states.** The four state variables are initialized as follows for each lesion $j$ of patient $i$:

- $T_j(0) = T_{ij1}$ — set to the first imaging observation for lesion $j$;
- $L(0) = 0$ — *CISH-KO* T-cells are supplied exclusively through the dose compartment;
- $N(0) = 0.1$ mm$^2$ — a small baseline level reflecting the depleted endogenous T-cell pool following lymphodepletion prior to infusion;
- $D_i(0) = N_{\text{infused}}/10^{10}$ — the scaled infused cell dose as described above.

**Parameter selection.** Model parameters are summarized in Table 2. The patient-level parameter set was determined using a recurrent elimination procedure guided by practical identifiability and cross-parameter correlation diagnostics. Starting from a broad candidate set, we iteratively removed parameters that the clinical data could not resolve independently, while retaining those that produced distinguishable signatures in the observed tumor and circulating ratio trajectories. Parameters related to endogenous T-cells ($k_N$, $d_N$, $\mu_N$, $\delta$, $h$) were therefore informed by prior studies or our mouse model fitting results and held at or near their previously estimated values with narrow ranges when they proved non-identifiable in the clinical fit. In contrast, *CISH-KO*-specific parameters ($k_L$, $d_L$, $\mu_L$, $p_L$, $f_{\text{dose}}$, $\kappa$) have not been characterized in previous studies; these were assigned broader feasible ranges and estimated from the clinical data. The tumor intrinsic growth rate $\gamma$ was allowed to vary across individual lesions within the range $[0.001, 0.1]$ day$^{-1}$, corresponding to tumor doubling times of approximately one to ten weeks, consistent with metastatic gastrointestinal cancers.

#### S2.3 Fitting Procedure

**Hierarchical (bi-level) optimization.** The calibration was performed using a nested two-level optimization scheme:

- *Outer level (patient-level).* For each candidate set of patient-level parameters, the algorithm solves the inner-level optimization for every tumor lesion of that patient and aggregates the per-tumor objectives into a single patient-level cost.
- *Inner level (tumor-level).* Given a fixed set of patient-level parameters, each tumor lesion $j$ is fitted independently for the tumor-specific parameter(s). In the present formulation the only tumor-level parameter is the intrinsic growth rate $\gamma_j$. Because this is a one-dimensional optimization, we employ Brent's method [13], a derivative-free bracketing algorithm, on the

| Par. | Unit | Description | Mouse 2015 | Mouse 2022 | Patient | Ref. |
|---|---|---|---|---|---|---|
| $\gamma$ | day$^{-1}$ | Tumor intrinsic growth rate | 0.2180 | 0.1780 | 0.01–0.1 | – |
| $K$ | mm$^2$ | Carrying capacity for total T-cell population | 1 | 1 | 1 | – |
| $k_L$ | day$^{-1}$ | Tumor killing rate by *CISH-KO* T-cells ($L$) | 0.5519 | 1.1048 | $10^{-3}$–1 | – |
| $k_{L_o}$ | day$^{-1}$ | Tumor killing rate by wild-type T-cells ($L_o$) | 0.4834 | 0.4834 | – | – |
| $k_N$ | day$^{-1}$ | Tumor killing rate by endogenous T-cells ($N$) | $10^{-4}$ | $10^{-4}$ | $5 \times 10^{-4}$ | – |
| $\mu_L$ | day$^{-1}$ mm$^{-2}$ | Tumor-induced exhaustion rate of $L$ | $1.2 \times 10^{-4}$ | $1.2 \times 10^{-4}$ | $4 \times 10^{-4}$ | – |
| $\mu_N$ | day$^{-1}$ mm$^{-2}$ | Tumor-induced exhaustion rate of $N$ | $4 \times 10^{-4}$ | $4 \times 10^{-4}$ | $4 \times 10^{-4}$ | [9] |
| $d_L$ | day$^{-1}$ | Natural death rate of $L$ | 0.05 | 0.3541 | $10^{-3}$–0.5 | – |
| $d_N$ | day$^{-1}$ | Natural death rate of $N$ | $3 \times 10^{-2}$ | $3 \times 10^{-2}$ | $3 \times 10^{-2}$ | [9] |
| $p_L$ | day$^{-1}$ | Proliferation rate of $L$ | 0.2642 | 0.3539 | $10^{-3}$–1 | – |
| $p_N$ | day$^{-1}$ | Proliferation rate of $N$ | $5 \times 10^{-2}$ | $5 \times 10^{-2}$ | $10^{-3}$–1 | [9] |
| $\delta$ | day$^{-1}$ | Tumor-induced recruitment rate of $N$ | 0.01 | 0.01 | 0.01 | [10] |
| $h$ | mm$^2$ | Half-saturation constant for endogenous T-cell recruitment | $10^3$ | $10^3$ | $10^3$ | [2] |
| $f_{\text{dose}}$ | day$^{-1}$ | Clearance rate of infused T-cells prior to engraftment | 0 | 0 | 0.01–1.0 | – |
| $\kappa$ | day$^{-1}$ | Rate of infiltration from dose compartment to $L$ | 0.1516 | 0.0446 | 0.05–0.5 | – |
| $\mu_Z$ | day$^{-1}$ | Natural decay rate of anti-PD-1 drug ($Z$) | – | 0.7 | – | – |
| $a$ | – | Shape parameter for $\alpha(Z)$ | – | 0.3815 | – | – |
| $b$ | – | Sigmoid steepness parameter for $\alpha(Z)$ | – | 50 | – | – |
| $b_L$ | – | Sigmoid steepness parameter for $\alpha_L(Z)$ | – | 0.01 | – | – |

Table 2: Model parameters, units, and interpretations for mouse experiments and clinical trials.

interval $[\gamma_{\text{lb}}, \gamma_{\text{ub}}] = [0.001, 0.1]$ day$^{-1}$, which is substantially faster than gradient-based methods for scalar problems.

**Objective function.** The total objective for patient $i$ is

$$\mathcal{L}^{(i)}_{\text{patient}} = \sum_{j=1}^{J_i} \mathcal{L}^{(ij)}_{\text{tumor}} + \mathcal{L}^{(i)}_{\text{CISH}}. \tag{7}$$

*Tumor size component:* For lesion $j$ of patient $i$ with observed data $\{(t_{ik}, T_{ijk})\}_{k=1}^{K_{ij}}$:

$$\mathcal{L}^{(ij)}_{\text{tumor}} = \sum_{k=1}^{K_{ij}} \left(1 - \frac{T_j^{\text{model}}(t_{ik})}{T_{ijk}}\right)^2, \tag{8}$$

where $T_j^{\text{model}}(t_{ik})$ is the ODE-predicted tumor area for lesion $j$ evaluated at the observation time $t_{ik}$.

*CISH-KO ratio component:* The model-predicted *CISH-KO*/wild-type ratio is averaged across all lesions and compared to the peripheral blood measurement $R_{ik}$:

$$\mathcal{L}_{\text{CISH}}^{(i)} = w \sum_{k=1}^{K_i^{\text{CISH}}} \left( \frac{R_{ik} - \hat{R}_i(s_{ik})}{R_{i1}} \right)^2, \tag{9}$$

where $w = 0.2$ is a weighting factor chosen to down-weight the ratio component relative to the tumor size fits, mitigating the influence of the blood-to-tumor ratio approximation on the overall parameter estimates (see Section S2.1),

$$\hat{R}_i(t) = \frac{\bar{L}_i(t)}{\bar{N}_i(t)}, \qquad \bar{L}_i(t) = \frac{1}{J_i} \sum_{j=1}^{J_i} L_j(t), \qquad \bar{N}_i(t) = \frac{1}{J_i} \sum_{j=1}^{J_i} N_j(t)$$

denote the tumor-averaged model-predicted ratio and cell populations, and $R_{i1}$ is the first post-infusion *CISH-KO* ratio observation for patient $i$.

All residuals in (9) are normalized by the first-time-point measurement $R_{i1}$ rather than the contemporaneous value $R_{ik}$. This choice reflects the biological observation that $R_{ik} \ll R_{i1}$ for all $k > 1$: the *CISH-KO*/wild-type ratio peaks shortly after infusion and decays rapidly as *CISH-KO* T-cells are cleared. At later time points the measured ratio becomes very small and approaches the detection floor of the assay, whereas the ODE-predicted ratio $\hat{R}_i(s_{ik})$ can decrease without bound. Normalizing by $R_{i1}$ therefore places the primary fitting emphasis on the early, high-signal time point while tolerating larger relative deviations at later times when both the data and the model output are near zero and the signal-to-noise ratio is inherently low.

**Parameter transformation.** To improve optimizer performance across parameters spanning multiple orders of magnitude, patient-level parameters were optimized in a power-law-transformed space. Given initial parameter values $\theta_0$, the optimization was conducted over transformed variables $b$ such that

$$\theta = b^{\alpha}, \qquad \alpha = \log_{10}(\theta_0). \tag{10}$$

The bounds on $\theta$ were correspondingly transformed, and the lower and upper bounds were rearranged to ensure $b_{\text{lb}} < b_{\text{ub}}$ in all dimensions.

**Three-stage optimization.** The outer (patient-level) optimization employed a three-stage procedure to balance broad exploration with local refinement.

1. **Stage 1 — Broad multistart with Latin Hypercube Sampling.** $N_{\text{start}} = 200$ initial points were generated via Latin Hypercube Sampling (LHS) over the transformed bound box $[b_{\text{lb}}, b_{\text{ub}}]$ to ensure uniform coverage of the parameter space. Each point was optimized using the L-BFGS algorithm with box constraints (Fminbox) and finite-difference gradient approximations, with relative function tolerance $10^{-6}$, up to 8 outer iterations, and 200 inner iterations per outer iteration. All starts were distributed across available CPU threads using Julia's multi-threading facility.

2. **Stage 2 — Local refinement of top candidates.** The $N_{\text{refine}} = 10$ solutions with the lowest objective values from Stage 1 were selected for further refinement. Each was re-optimized with L-BFGS at tighter tolerances (relative function tolerance $10^{-8}$, up to 10 outer iterations, 300 inner iterations), followed by a Nelder–Mead simplex search (relative function tolerance $10^{-8}$, up to 500 iterations) as a derivative-free polish to escape potential gradient-based traps. The Nelder–Mead result was accepted only if it remained within the feasible region and improved the objective.

3. **Stage 3 — Final polish of best solution.** The single best solution from Stages 1–2 was subjected to a final round of L-BFGS optimization at very tight tolerances (relative function tolerance $10^{-10}$, gradient tolerance $10^{-10}$, 12 outer iterations, 500 inner iterations), followed again by a Nelder–Mead polish ($10^{-10}$ relative tolerance, 500 iterations).

The parameter set yielding the lowest overall objective across all three stages was selected as patient $i$'s best-fit solution.

**Numerical solver.** The ODE system (6) was integrated using the Tsitouras 5th-order Runge–Kutta method (Tsit5) implemented in `DifferentialEquations.jl`, with absolute tolerance $10^{-10}$, relative tolerance $10^{-3}$, and the solution saved at 0.5-day intervals over each patient's observation window.

**Software.** All fitting was implemented in Julia (v1.10+) using `DifferentialEquations.jl` for ODE integration, `Optim.jl` for optimization, and `StaticArrays.jl` for type-stable fixed-size arrays. Source code is available at Github repository to be created soon.

#### S2.4 Parameter Identifiability Analysis

The patient-level parameter set was determined through an iterative elimination procedure informed by practical identifiability and cross-patient correlation diagnostics. Consistent with the reduce-from-complete modeling strategy described in the main text, we began with the full set of biologically motivated candidate parameters and systematically removed those that could not be independently resolved from the available clinical data, rather than restricting the parameter set *a priori*. This data-driven reduction ensures that the final parameterization reflects the information content of the clinical measurements, and that the retained parameters correspond to mechanistically distinct processes whose individual contributions are distinguishable in the observed tumor and T-cell dynamics.

**Correlation analysis.** After fitting all 12 patients, we computed the Pearson correlation matrix of the optimal parameter values across patients (treating each patient's estimate $\hat{\theta}_i$ as one observation). Parameter pairs with $|\rho| > 0.6$ were flagged as potentially redundant, indicating that the two parameters may compensate for each other.

**Profile likelihood analysis.** To assess practical identifiability, we performed a profile likelihood analysis for every patient-level parameter. For parameter $\theta_m$ and patient $i$, $\theta_m$ was fixed at each of seven multiplier values $\alpha \in \{0.7, 0.8, 0.9, 1.0, 1.1, 1.2, 1.3\}$ times its optimal estimate $\hat{\theta}_{i,m}$; for each fixed value the remaining parameters were re-optimized, producing a profile $\mathcal{L}_i(\alpha;\, \theta_m)$.

Because a parameter may be identifiable in some patients but not others (the profile is a local analysis centered at each patient's optimal value), we aggregated across all patients by normalizing each per-patient profile by its value at the optimum ($\alpha = 1$) and summing:

$$\mathcal{P}(\alpha;\, \theta_m) = \sum_{i=1}^{N_p} \frac{\mathcal{L}_i(\alpha;\, \theta_m)}{\mathcal{L}_i(1;\, \theta_m)}. \tag{11}$$

A parameter was considered practically identifiable if $\mathcal{P}$ exhibited a clear minimum with well-defined curvature at $\alpha = 1$; a flat profile ($\mathcal{P} \approx N_p$ for all $\alpha$) indicated practical non-identifiability.

**Iterative elimination.**

1. **Round 1.** We began with the full set of 11 candidate patient-level parameters:

   $$\Theta_1 = \{d_L,\, k_L,\, p_L,\, \mu_L,\, d_N,\, k_N,\, p_N,\, \mu_N,\, \delta,\, f_{\text{dose}},\, \kappa\}.$$

   The model was fitted to all 12 patients and the diagnostics were computed.

   *Correlation* (Figure 1): Several parameter pairs exhibited high correlations, notably ($\mu_N$, $\kappa$) with $\rho = 0.64$ and ($k_L$, $\mu_N$) with $\rho = 0.62$.

   *Profile likelihood* (Figure 2a and Figure 2b): $\mu_N$ and $\delta$ showed essentially flat aggregate profiles ($\mathcal{P}$ varied by less than $10^{-4}$ across all multipliers), indicating that these parameters are practically non-identifiable from the available data. Furthermore, $\mu_N$ was highly correlated with $k_L$. Both $\mu_N$ and $\delta$ were therefore removed.

   Biologically, the non-identifiability of $\delta$ reflects the fact that tumor-stimulated recruitment of endogenous T-cells cannot be resolved independently from their intrinsic proliferative expansion ($p_N$) given the available clinical data: both mechanisms produce the same qualitative effect—an increase in endogenous T-cell abundance in response to tumor burden—and the longitudinal tumor size and circulating ratio measurements do not provide sufficient information to disentangle the two. The non-identifiability of $\mu_N$, and its high correlation with $k_L$, arises because tumor-induced exhaustion of endogenous T-cells and *CISH-KO* T-cell-mediated tumor killing produce partially compensatory effects on the observed tumor trajectory: a higher exhaustion rate reduces the endogenous T-cell population, which in turn reduces competition for the *CISH-KO* population and can mimic the effect of a higher $k_L$. Because the data primarily constrain the net rate of tumor size change rather than the individual T-cell population dynamics, these two parameters trade off against each other.

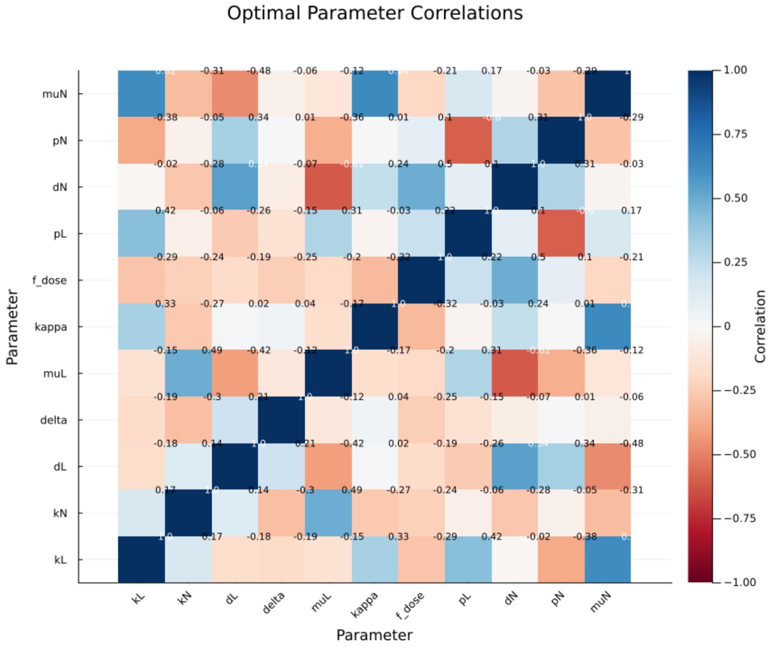


Figure 1: Pearson correlation heatmap of optimal patient-level parameters after Round 1 fitting ($|\Theta_1| = 11$).

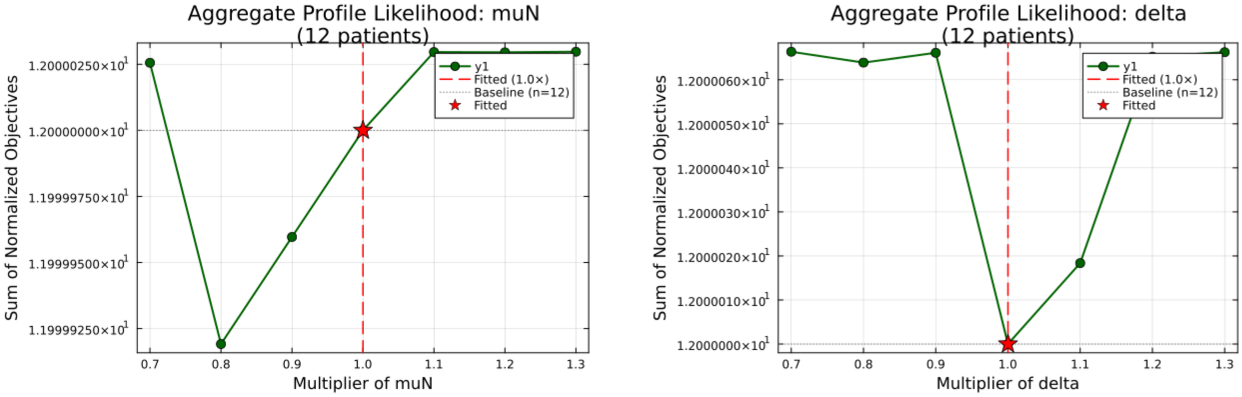


(a) $\mu_N$

(b) $\delta$

Figure 2: Aggregate profile likelihood for the two parameters removed after Round 1. Both profiles are essentially flat (considering the scaling), indicating practical non-identifiability.

2. **Round 2.** The reduced 9-parameter set was re-fitted:

$$\Theta_2 = \{d_L,\, k_L,\, p_L,\, \mu_L,\, d_N,\, k_N,\, p_N,\, f_{\text{dose}},\, \kappa\}.$$

*Correlation* (Figure 3): The pair $(p_L,\, d_N)$ exhibited $\rho = 0.97$, and $(k_L,\, k_N)$ had $\rho = -0.61$.

*Profile likelihood* (Figure 4a and Figure 4b): $d_N$ and $k_N$ showed weak profiles with limited curvature, while their correlated partners $p_L$ and $k_L$ exhibited substantially sharper minima. $d_N$ and $k_N$ were therefore removed.

The non-identifiability of $d_N$ and its near-perfect correlation with $p_L$ ($\rho = 0.97$) reflects a fundamental tradeoff: faster natural turnover of endogenous T-cells ($d_N$) frees niche space within the shared carrying capacity, which has a similar effect on the model-predicted tumor trajectory as increased proliferation of *CISH-KO* T-cells ($p_L$), since both mechanisms alter the competitive balance between the two populations. The weak identifiability of $k_N$ arises because endogenous T-cells have substantially lower cytotoxic capacity than *CISH-KO* T-cells ($k_N \ll k_L$), so moderate perturbations in $k_N$ produce only small changes in the overall rate of tumor killing; the tumor size data are therefore primarily informative about $k_L$, which dominates the cytotoxic signal.

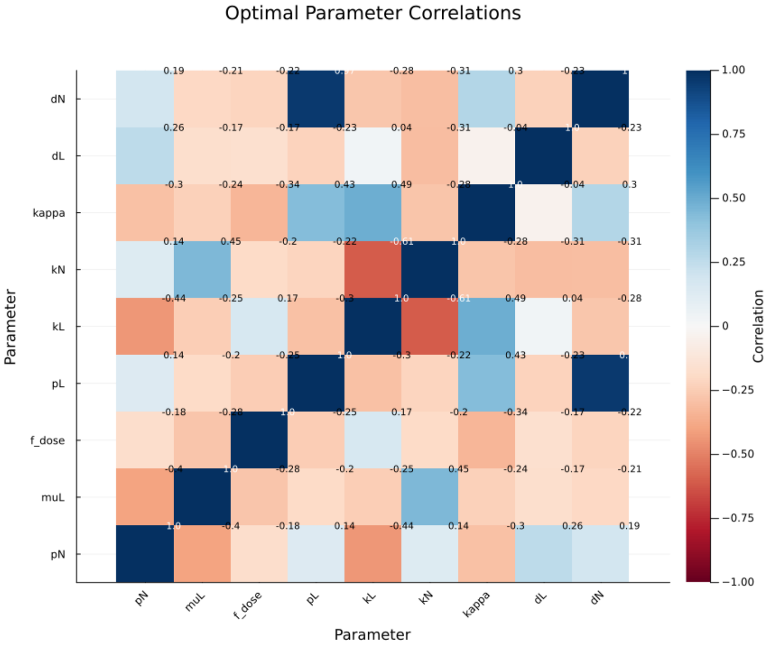


Figure 3: Pearson correlation heatmap of optimal patient-level parameters after Round 2 fitting ($|\Theta_2| = 9$).

3. **Round 3 (final).** The 7-parameter set was re-fitted and re-assessed:

$$\Theta_3 = \{d_L,\, k_L,\, p_L,\, \mu_L,\, p_N,\, f_{\text{dose}},\, \kappa\}.$$

All remaining parameters showed acceptable identifiability in the aggregate profile likelihood (Table 3; individual profiles in Figures 6a–6g) and pairwise correlations with $|\rho| < 0.72$ (Figure 5). This set was adopted as the final patient-level parameter set. We note that further removal of $p_N$ or $d_L$ was tested but resulted in a substantial degradation of fitting

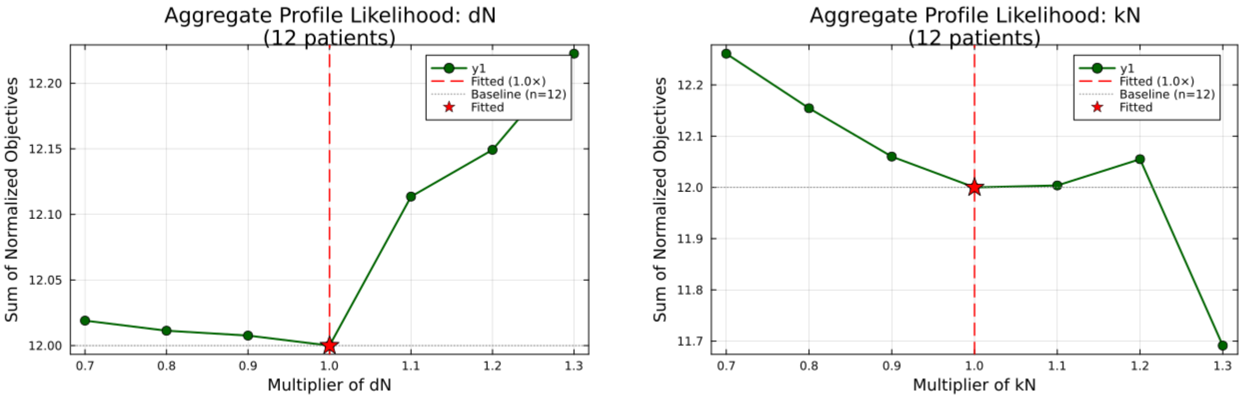


(a) $d_N$

(b) $k_N$

Figure 4: Aggregate profile likelihood for the two parameters removed after Round 2. Both profiles show weak curvature compared to their correlated partners ($p_L$ and $k_L$, respectively).

performance across patients, confirming that these parameters are necessary for the model to capture the data. This is consistent with the biological analysis in the main text: $d_L$ and $p_N$ emerge as dominant determinants of therapeutic outcome (Figure 5b,c), and their identifiability reflects the fact that they govern mechanistically distinct processes—*CISH-KO* T-cell survival and endogenous immune reconstitution, respectively—that are not confounded by other model parameters within the resolution of the available data.

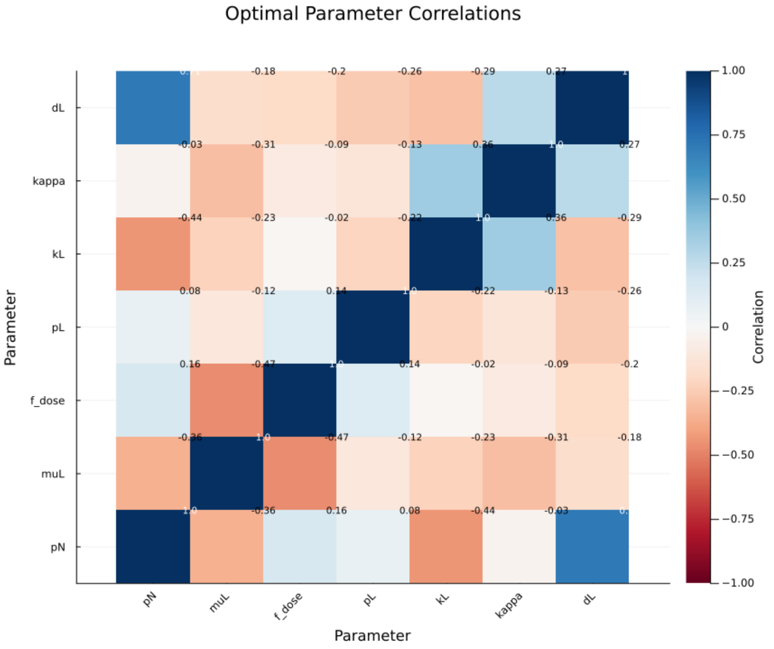


Figure 5: Pearson correlation heatmap of optimal patient-level parameters after Round 3 fitting ($|\Theta_3| = 7$). All pairwise correlations satisfy $|\rho| < 0.72$.

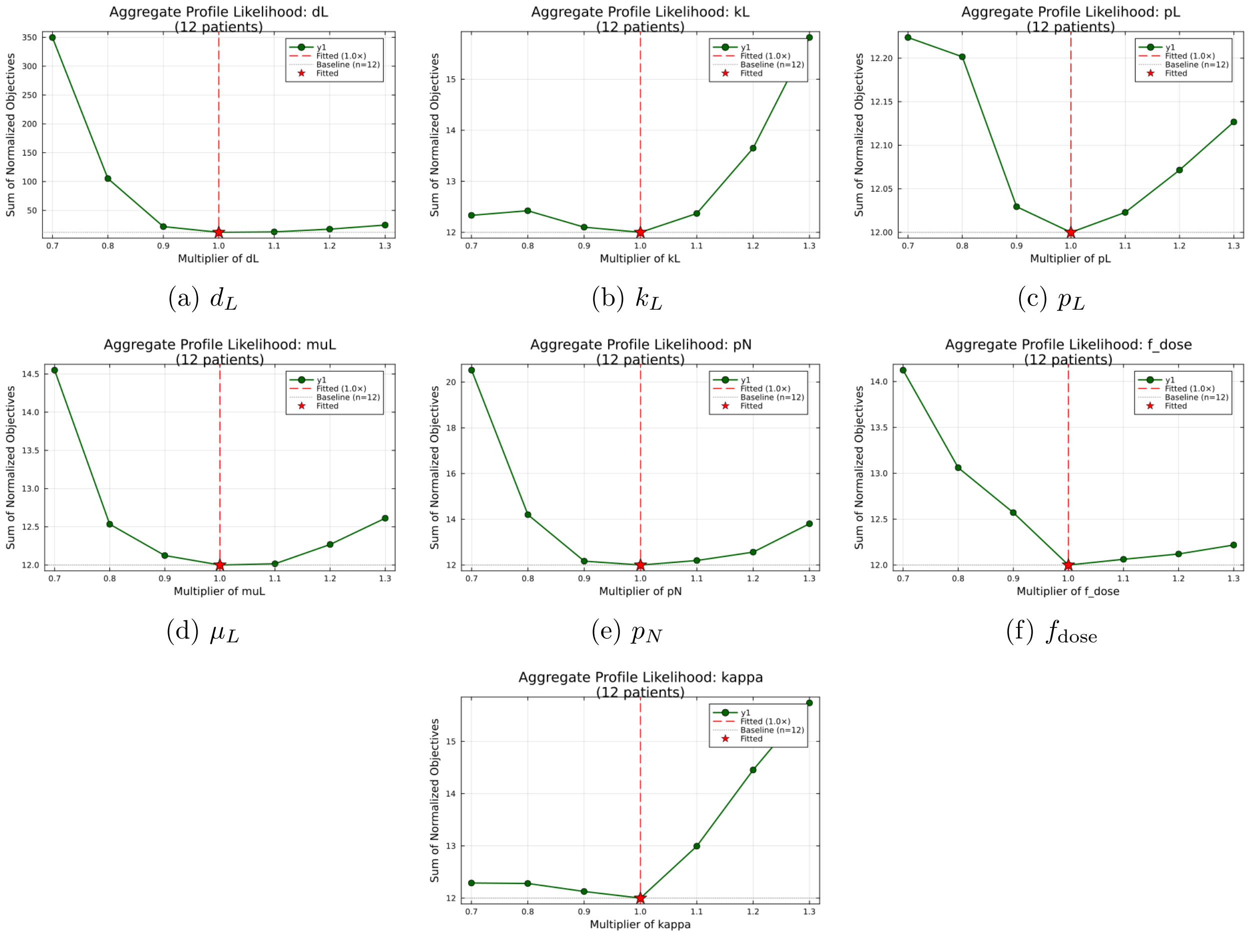


(a) $d_L$ (b) $k_L$ (c) $p_L$

(d) $\mu_L$ (e) $p_N$ (f) $f_{\text{dose}}$

(g) $\kappa$

Figure 6: Aggregate profile likelihood for all seven retained parameters after Round 3. Each profile exhibits a clear minimum at the fitted value ($\alpha = 1.0$), confirming practical identifiability.

The parameters removed through this process ($\mu_N$, $\delta$, $d_N$, $k_N$) were fixed at their nominal values informed by prior mouse model experiments (Table 2). Collectively, the non-identifiable parameters share a common feature: they govern aspects of endogenous T-cell dynamics—recruitment ($\delta$), exhaustion ($\mu_N$), natural death ($d_N$), and cytotoxic capacity ($k_N$)—that are not directly observed in the clinical data. Because the available measurements constrain the system primarily through longitudinal tumor sizes and the circulating *CISH-KO*/endogenous T-cell ratio, endogenous T-cell dynamics are observed only indirectly through their aggregate effect on tumor growth and competitive pressure on the *CISH-KO* population. This renders individual endogenous T-cell kinetic parameters difficult to resolve, while the net competitive effect of the endogenous compartment remains well captured by $p_N$, which controls the rate at which endogenous T-cells reconstitute and occupy the shared immune niche.

| Parameter | Round 1 | Round 2 | Round 3 | Status |
|---|---|---|---|---|
| $\kappa$ | 13.98 | 14.34 | 12.13 | Retained |
| $d_L$ | 13.32 | 12.40 | 21.94 | Retained |
| $k_L$ | 12.07 | 12.08 | 12.10 | Retained |
| $p_N$ | 12.64 | 13.54 | 12.17 | Retained |
| $f_{\text{dose}}$ | 12.39 | 12.42 | 12.57 | Retained |
| $\mu_L$ | 12.04 | 12.30 | 12.12 | Retained |
| $p_L$ | 12.00 | 11.82 | 12.03 | Retained |
| $d_N$ | 12.00 | 12.01 | | Removed (Round 2) |
| $k_N$ | 12.03 | 12.06 | | Removed (Round 2) |
| $\delta$ | 12.00 | | | Removed (Round 1) |
| $\mu_N$ | 12.00 | | | Removed (Round 1) |

Table 3: Aggregate profile likelihood curvature across elimination rounds. Values report $\mathcal{P}(0.9;\theta_m)$, the aggregate normalized objective at the 0.9× multiplier (with $N_p = 12$ patients, so a baseline of 12.0). Higher values indicate sharper profiles and better identifiability.

#### S2.5 Individual Fitting Results

Patient-level fitted parameters are reported in Table 4, and individual patient fits are shown in Figures 7–9.

| Patient | $\gamma$ | $d_L$ | $k_L$ | $\mu_L$ | $p_L$ | $p_N$ | $f_{\text{dose}}$ | $\kappa$ |
|---|---|---|---|---|---|---|---|---|
| UMN 002 | 0.024104, 0.007168, 0.014848, 0.014269 | 0.0197 | 0.0365 | $1.19 \times 10^{-8}$ | 0.0011 | 0.1539 | 0.0102 | 0.5 |
| UMN 003 | 0.028905, 0.024713 | 0.0307 | 0.0302 | $8.73 \times 10^{-7}$ | 0.001 | 0.2288 | 1 | 0.0666 |
| UMN 006 | 0.004699 | 0.0011 | 0.001 | $1.04 \times 10^{-8}$ | 0.2257 | 0.4997 | 0.7524 | 0.1181 |
| UMN 009 | 0.055991, 0.046328, 0.026503, 0.018719 | 0.4202 | 0.0064 | $6.75 \times 10^{-9}$ | 0.001 | 0.4717 | 0.2933 | 0.5 |
| UMN 012 | 0.009089, 0.007057, 0.009042, 0.009126, 0.010052 | 0.003 | 0.0025 | 0.7728 | 0.0029 | 0.0664 | 0.0174 | 0.05 |
| UMN 014 | 0.023525, 0.023217, 0.016795, 0.096173, 0.099999 | 0.1637 | 0.0066 | $1.41 \times 10^{-8}$ | 0.001 | 0.3686 | 1 | 0.4395 |
| UMN 015 | 0.031886, 0.026031 | 0.0591 | 0.0113 | $1.01 \times 10^{-9}$ | 0.001 | 0.3747 | 1 | 0.05 |
| UMN 017 | 0.021546, 0.020352, 0.064593, 0.060318 | 0.2555 | 0.0024 | $5.02 \times 10^{-8}$ | 0.001 | 1 | 1 | 0.2327 |
| UMN 018 | 0.052225, 0.046634 | 0.5 | 0.001 | 0.1109 | 0.001 | 0.8874 | 0.01 | 0.05 |
| UMN 019 | 0.013866, 0.01306, 0.010234, 0.029183 | 0.0352 | 0.001 | $2.45 \times 10^{-8}$ | 0.0016 | 0.3221 | 1 | 0.05 |
| UMN 020 | 0.015923, 0.022676 | 0.0288 | 0.0014 | 0.0798 | 0.0011 | 0.3213 | 0.011 | 0.05 |
| UMN 022 | 0.001 | 0.001 | 0.7661 | $6.62 \times 10^{-4}$ | 0.001 | 0.3088 | 1 | 0.05 |

Table 4: Patient-level fitted parameters. For patients with multiple tumor lesions, the tumor growth rate $\gamma$ is reported separately for each lesion, while all other parameters are shared at the patient level.

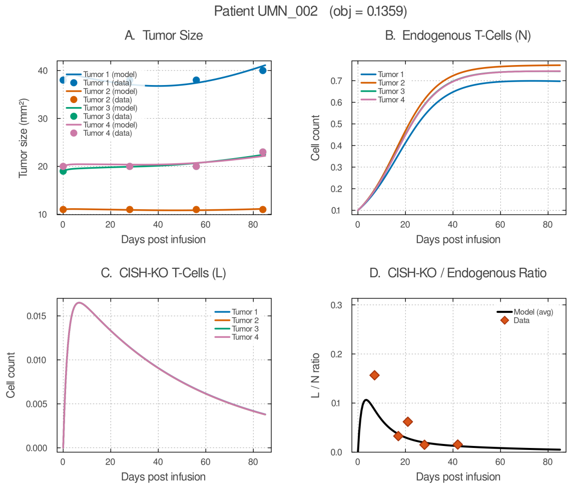


(a) Patient UMN_002

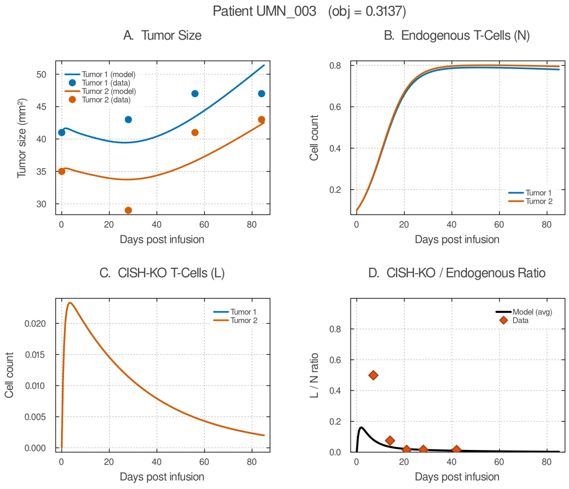


(b) Patient UMN_003

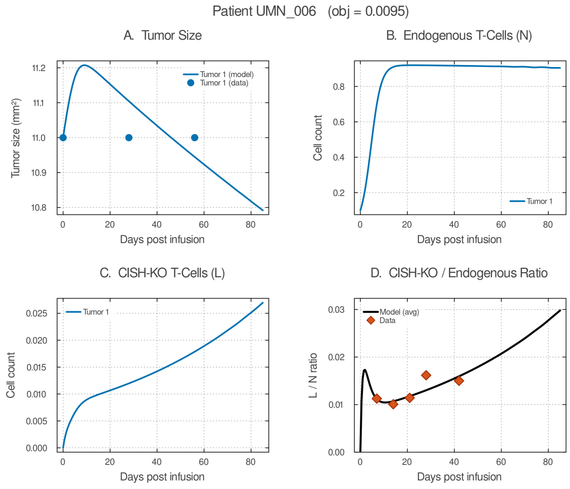


(c) Patient UMN_006

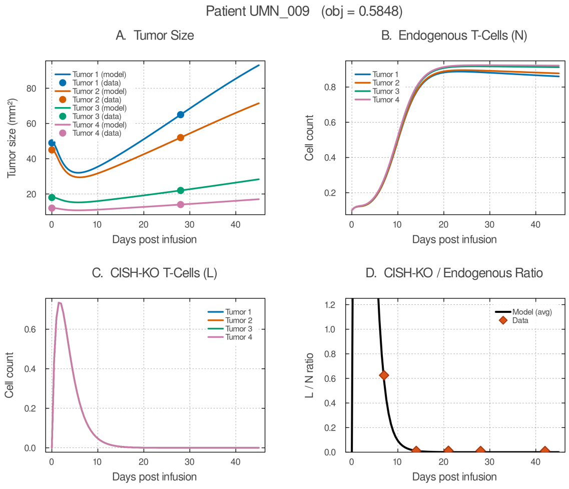


(d) Patient UMN_009

Figure 7: Individual patient model fits (patients UMN_002–UMN_009). Each sub-panel shows **(A)** tumor size, **(B)** endogenous T cells, **(C)** *CISH-KO* cells, and **(D)** *CISH-KO*/endogenous ratio. Solid lines: model; markers: data.

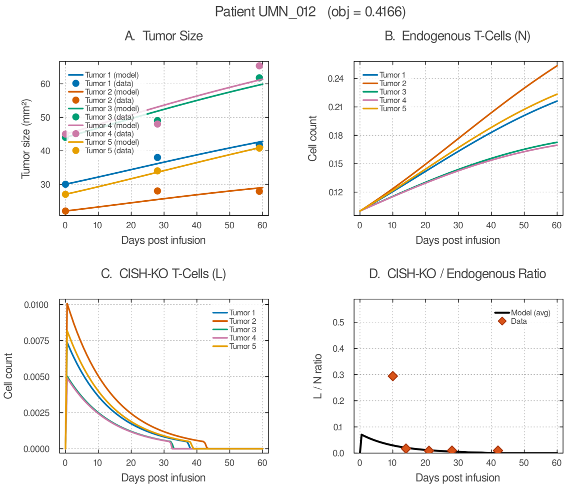


(a) Patient UMN_012

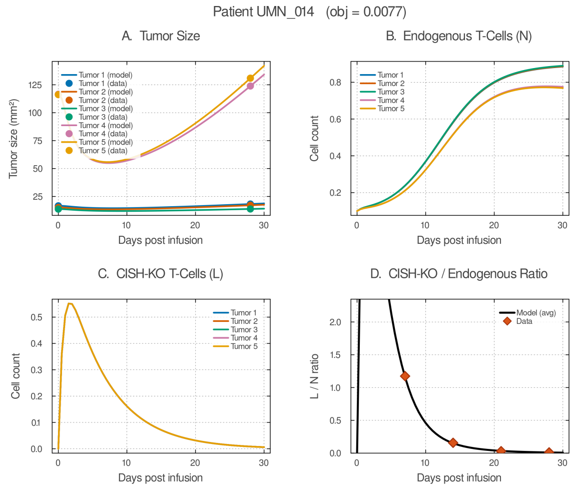


(b) Patient UMN_014

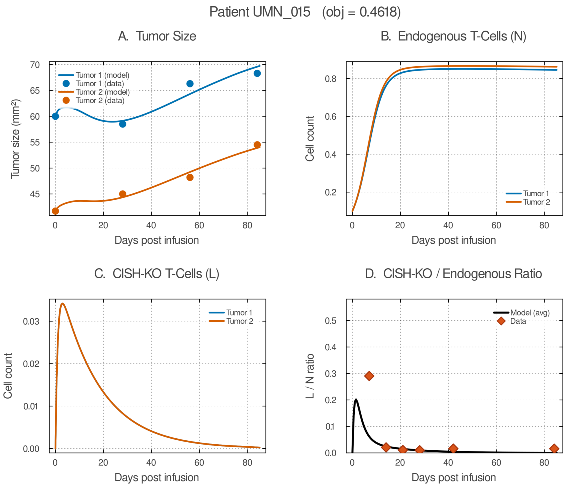


(c) Patient UMN_015

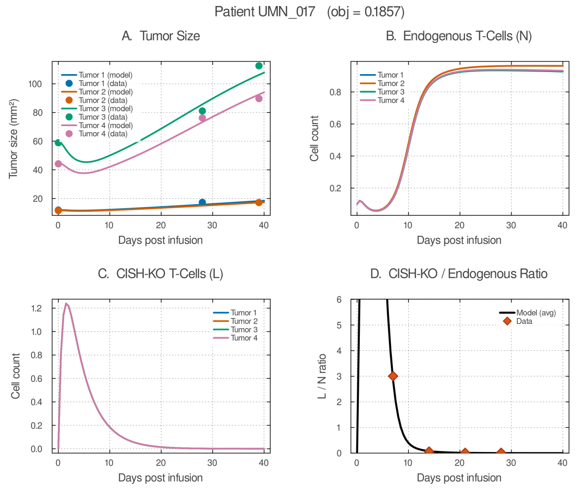


(d) Patient UMN_017

Figure 8: Individual patient model fits (patients UMN_012–UMN_017). Layout as in Fig. 7.

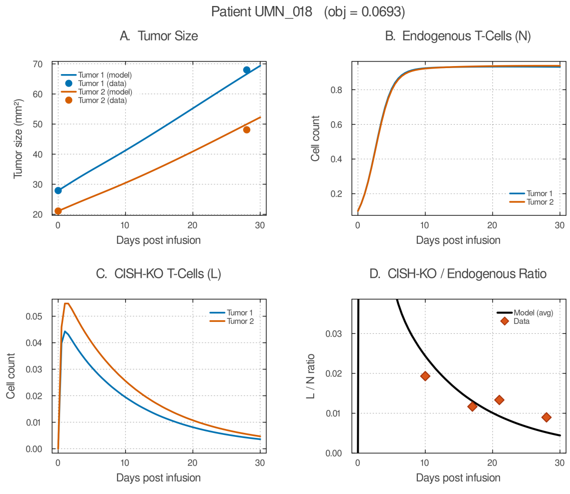


(a) Patient UMN_018

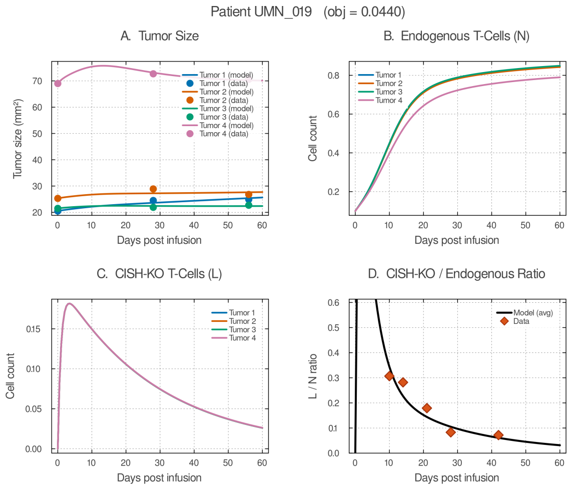


(b) Patient UMN_019

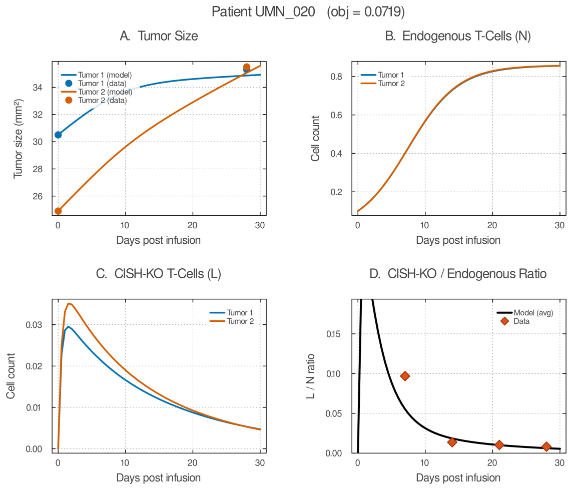


(c) Patient UMN_020

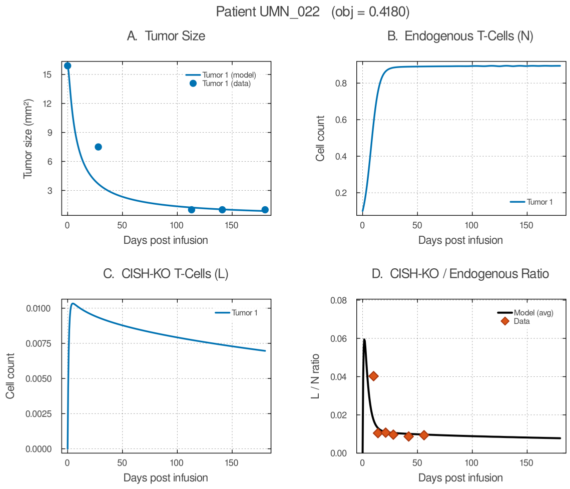


(d) Patient UMN_022

Figure 9: Individual patient model fits (patients UMN_018–UMN_022). Layout as in Fig. 7.
